# KDM6A loss sensitizes human acute myeloid leukemia to PARP and BCL2 inhibition

**DOI:** 10.1101/2022.07.12.498585

**Authors:** Liberalis Debraj Boila, Subhadeep Ghosh, Subham K. Bandyopadhyay, Liqing Jin, Alex Murison, Andy G. X. Zeng, Wasim Shaikh, Satyaki Bhowmik, Siva Sai Naga Anurag Muddineni, Mayukh Biswas, Sayantani Sinha, Shankha Subhra Chatterjee, Nathan Mbong, Olga I. Gan, Anwesha Bose, Sayan Chakraborty, Andrea Arruda, James A. Kennedy, Amanda Mitchell, Eric R. Lechman, Debasis Banerjee, Michael Milyavsky, Mark D. Minden, John E. Dick, Amitava Sengupta

## Abstract

Acute myeloid leukemia (AML) is a heterogeneous, aggressive malignancy with dismal prognosis and with limited availability of targeted therapies. AML exhibits epigenetic deregulation and transcriptional plasticity that contributes to pathogenesis. KDM6 proteins are histone-3 lysine-27 demethylases that play major context dependent roles in AML evolution and therapy resistance. Here, we demonstrate that KDM6 demethylase function critically regulates DNA damage repair (DDR) gene expression programs in AML. Mechanistically, KDM6 family protein expression is regulated by genotoxic stress, with deficiency of KDM6A (UTX) and KDM6B (JMJD3) impairing DDR transcriptional activation and compromising repair potential. Acquired KDM6A *loss-of-function* mutations have been implicated in chemoresistance, although a significant percentage of relapsed AML have upregulated KDM6A. Based on these mechanistic findings, olaparib treatment significantly reduced engraftment of patient-derived xenografts. Thus *KDM6A*-mutant human primary AML samples have increased susceptibility to Poly-(ADP-ribose)-polymerase (PARP) inhibition *in vivo*. Crucially, a higher KDM6A expression is correlated with venetoclax tolerance. Loss of KDM6A increased mitochondrial activity, BCL2 expression, and sensitized AML cells to venetoclax. Additionally, KDM6A loss was accompanied with a downregulated BCL2A1, which is commonly associated with venetoclax resistance. Corroborating these results, dual targeting of PARP and BCL2 was superior to PARP or BCL2 inhibitor monotherapy in inducing AML apoptosis, and primary AML cells carrying acquired *KDM6A*-domain mutations were even more sensitive to the combination. Together, our study illustrates a mechanistic rationale in support for a novel combination targeted therapy for human AML based on subtype heterogeneity, and establishes KDM6A as an important molecular regulator for determining therapeutic efficacy.

## INTRODUCTION

KDM6 proteins represent a family of histone lysine demethylases that play an important role in chromatin remodeling and transcriptional regulation during multi-cellular development, establishment of tissue identity and tumorigenesis (*1–4*). KDM6A (UTX) and KDM6B (JMJD3) critically regulate demethylation of H3K27 methyl residues, whereas the catalytic potential of KDM6C (UTY) is poorly understood (*1, 5–7*). Growing evidence suggests involvement of KDM6A in acute myeloid leukemia (AML) pathogenesis (*4, 8–12*). KDM6A escapes X chromosome inactivation, and *Utx* null homozygous female mice spontaneously develop aging associated myeloid leukemia (*9, 13*). In addition, *KDM6A loss of function* mutation is implicated in conventional chemotherapy relapse in AML, indicating tumor suppressor function (*8, 10, 14*). KDM6A condensation, which involves a core intrinsically disordered region (cIDR), has been reported to confer tumour-suppressive activity independent of Jumonji C (JmjC) demethylase function (*14*). Recent studies suggested downregulation of KDM6A expression occurs in about 46% of cytogenetically normal karyotype and AraC relapsed AML patients (*8*). However, 37% of cases exhibited upregulated KDM6A transcripts. Thus KDM6A must function in a highly contextual fashion since there are subsets of AML cases where expression is on opposite ends of a spectrum. Therefore, the cause and pathophysiological relevance of KDM6A upregulation at chemotherapy relapse, observed in more than a third of the patients, is an open question. Additionally, to what extent KDM6A expression and function are connected with AML targeted therapy is unknown.

By contrast, KDM6B predominantly plays a context-dependent oncogenic function in haematological malignancies (*15, 16*). KDM6B regulates transcriptional elongation, and KDM6B expression is upregulated in myelodysplastic syndromes-hematopoietic stem/progenitors (*17, 18*). While KDM6A acts as a tumor suppressor and is frequently mutated in T-ALL, KDM6B is essential for the initiation and maintenance of T-ALL (*19, 20*). However, a subgroup of T-ALL expressing TAL1 is uniquely vulnerable to KDM6A inhibition (*21*). Together, KDM6A and KDM6B possess cell type-specific functions in leukemia, with KDM6 proteins and their associated signaling emerging as important focal points for developing molecular targeted therapy. Key cellular processes impacted by KDM6 demethylases include Th-cell development, integrated stress response activation, and regulation of DNA double stranded break repair.

Efficient repair of DNA damage caused by genotoxic stress is important for tissue homeostasis (*22, 23*). Tumor cells accumulate considerable levels of DNA damage and require robust DNA damage repair (DDR) mechanisms for survival (*24, 25*). AML cell survival depends upon an intact DNA repair machinery, with accumulation of DNA double stranded breaks (DSBs) leading to apoptosis (*26*). DSBs are among the most lethal DNA aberrations, and are repaired through either homologous recombination (HR) mediated repair or non-homologous end joining (NHEJ) (*23*). Targeting DNA repair pathways for cancer therapy has gained a momentum over the past few years, with poly(adenosine 5’-diphosphate-ribose) polymerase (PARP) inhibition for HR deficient tumors have shown promise in clinical settings (*27–29*). Therefore, identifying molecular regulation of DNA repair pathways important for AML cell survival is essential for developing effective combination targeted therapy.

Here we demonstrate that KDM6 demethylases play an important role in DDR gene regulation in AML opening the potential for improved molecular targeted therapies in AML through epigenetic modulation. Together, our study addresses two important clinical questions: first, PARP inhibition would be effective for KDM6A deficient AML, and secondly, KDM6A inhibition should potentiate PARP or BCL2 blockade in distinct subtypes of AML where KDM6A expression is upregulated or even maintained above threshold level.

## RESULTS

### KDM6 demethylases associate with DSB repair gene expression in AML

Kdm6a deficient homozygous female mice *(Utx^-/-^)* spontaneously develop aging associated AML (*9*). To identify genes regulated by KDM6A in AML development, we re-analyzed the available RNA-seq results from *Utx^-/-^*female mice presenting with AML (ERS1090539, ERS1090541, ERS1090542), compared to *Utx^+/+^* control females (ERS539514, ERS539515) (*9*). *Utx^-/-^* and MLL-AF9 negative AML splenocytes were able to propagate leukemia in secondary recipients. Deficiency of Kdm6a led to 4014 genes being downregulated and 4703 genes upregulated (FDR: 0.01; Log_2_FC: > 1.5) (dataset S1). KDM6A JmjC-demethylase function is predominantly associated with transcriptional activation (*30*). Gene ontology (GO) enrichment analysis of the downregulated genes (4014) in *Utx^-/-^*cells revealed enrichment of several GO terms linked with DNA repair, with the most significant being the double strand break (DSB) repair (Fig. 1A). The DSB repair term included genes of both HR and NHEJ pathways (fig. S1A). Re-analysis of ChIP-seq results conducted in *Utx^+/+^* hematopoietic cells showed a total of 1825 Kdm6a ChIP-seq occupied genes (*9*), which were downregulated upon Kdm6a loss. GO analysis of these 1825 genes (dataset S2) further revealed a significant enrichment of DNA repair associated GO terms, suggesting involvement of Kdm6a demethylase in DNA repair (fig. S1B).

**Fig. 1.**
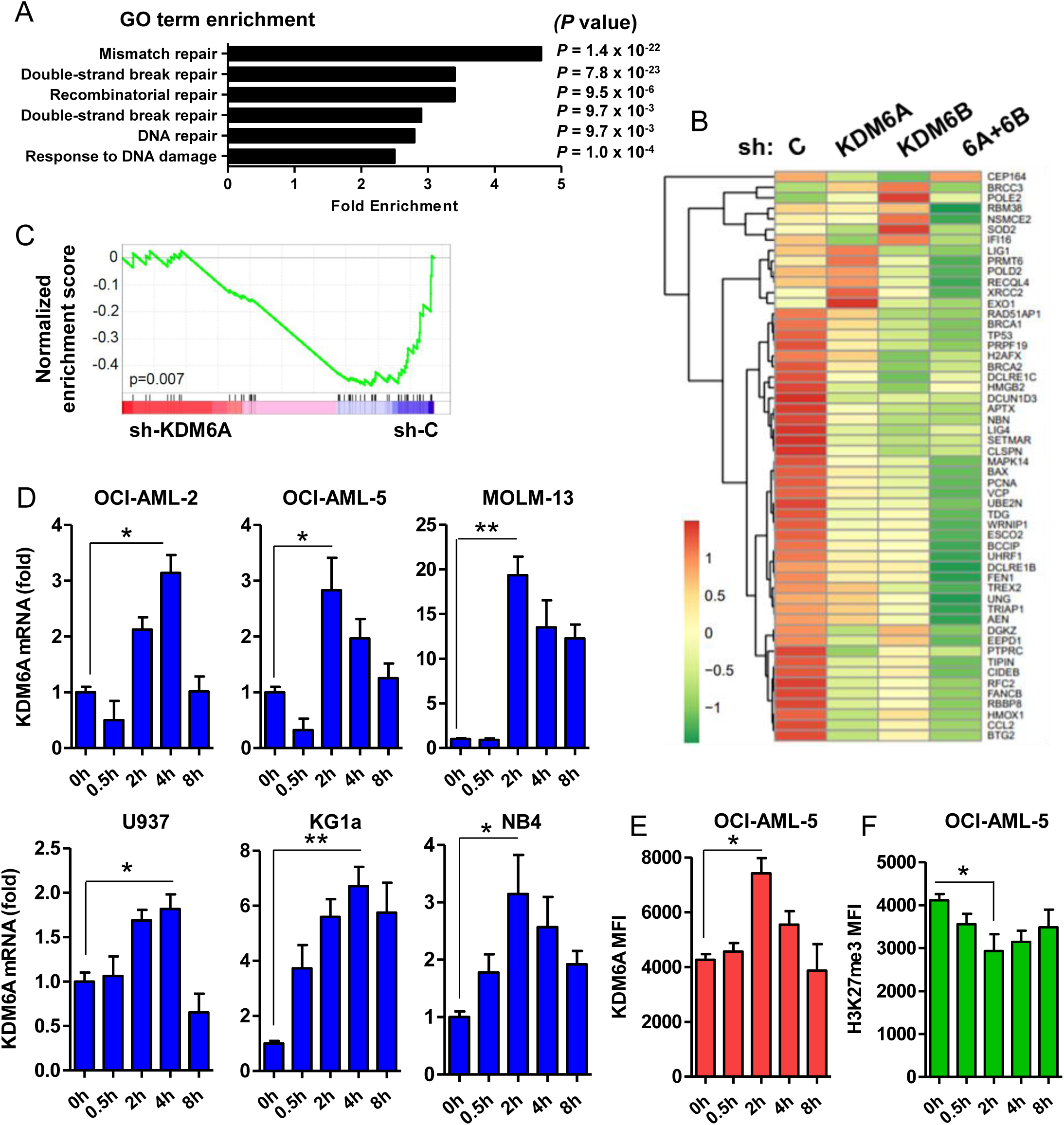
KDM6 demethylases associate with DNA repair gene expression in AML. (A) Gene Ontology (GO) term analysis of the 4014 downregulated genes in *Utx^-/-^* AML splenocytes. (B) RNA-seq heatmap showing expression of DDR genes in control and KDM6A and/or KDM6B deficient U937 cells (n=5). (C) Gene Set Enrichment Analysis (GSEA) for DDR pathway genes in KDM6A deficient U937 cells compared to control. (D) Quantitative reverse transcriptase-PCR (qRT-PCR) analysis (normalized to 0h) of KDM6A in AML cells treated with 10 Gy γ-irradiation (γ-IR) (n=3). (E) Flow cytometry analysis showing mean fluorescence intensity (MFI) of KDM6A in OCI-AML-5 cells irradiated with 10 Gy (n=3). (F) Flow cytometry analysis showing MFI of H3K27me3 in OCI-AML-5 cells irradiated with 10 Gy (n = 3). qRT-PCR values were normalized to GAPDH. Data are representative of at least three independent experiments. Statistics were calculated with Student’s t-test; error bars represent means ± SD. \**P* < 0.05 or \*\**P* < 0.01 were considered to be statistically significant.

To dissect the role of KDM6 proteins in regulating DDR gene expression in AML, we generated U937 cells using *shRNA-*expressing lentivirus vectors against KDM6A or KDM6B or both (hereafter referred as KDM6 deficient cells) (fig. S1C). U937 cell, originally isolated from a patient with histiocytic lymphoma, has been defined as a promonocytic myeloid leukemia cell line, capable of monocytic differentiation and has frequently been used as a model for myeloid leukemia. Additionally, U937 cells are relatively resistant to standard chemotherapy including KDM6 small molecule inhibitor GSK-J4, and therefore can serve as a relevant model to characterize targeted therapy. KDM6 knockdown led to an increase in global H3K27me3 and a decrease in H3K27ac levels, with the difference being more prominent in KDM6B knockdown and double knockdown cells (fig. S1D). Deficiency of KDM6A and/or KDM6B did not affect proliferation of U937 cells (fig. S1E). KDM6A deficient AML cell lines did not show consistent change in myeloid differentiation (fig. S1F). RNA-seq analysis (FDR: 0.05; Log_2_FC: > 2) suggested that there was significant downregulation (*P* < 0.05) of > 80 DDR genes upon knockdown of KDM6A alone or KDM6B alone or both (Fig. 1B). Gene Set Enrichment Analysis (GSEA) uncovered a significantly lower enrichment of DDR pathway in KDM6A deficient AML (Fig. 1C). GO term also indicated enrichment of multiple DNA repair genes, including BRCA and RAD families, in KDM6 deficient AML (fig. S1G). Together, these findings suggest that KDM6 proteins are associated with DDR gene regulation in AML.

### DSB repair activation induces expression of KDM6 in AML

To elucidate the function of KDM6A in mediating DSB repair in AML, we first interrogated irradiation induced alteration of KDM6A in AML cells. H3K27me3 level influences DSB repair efficiency, and decrease in H3K27me3 associates with radiation dosage, with 10 Gy irradiation causing maximum decrease (*31*). Interestingly, a single dose of γ-radiation (10 Gy) induced a time-dependent increase in expression of KDM6A (6 out of 6 AML cell lines tested) and KDM6B (4 out of 6 lines tested) independent of pathological or molecular subtypes (Fig. 1D and fig. S1H). Increase in KDM6A expression was observed as early as 30 min in KG1a cells, while OCI-AML-2 and KG1a cells showed maximum induction at 4 hours after irradiation (Fig. 1D). Low dose irradiation in AML cells did not sufficiently induce expression of KDM6A or KDM6B (fig. S2A). In agreement with gene expression alteration, KDM6A protein was also upregulated on radiation accompanied with a concomitant decrease in H3K27me3 (Figs. 1E-F and figs. S2B-C). There was no significant induction of KDM6A or KDM6B in normal CD34^+^CD38^-^CD45RA^-^ hematopoietic stem cells (HSCs) upon genotoxic stress (fig. S2D). Collectively, these results indicate that γ-IR mediated DNA repair induces KDM6 demethylase expression in AML.

### Deficiency of KDM6 impairs DDR gene expression and DSB repair in AML

Efficient DSB repair has been shown to promote survival of AML cells (*26*). To identify target genes that sensitize AML cells to genotoxic stress, we leveraged a previously reported genome-wide pooled lentiviral *shRNA* screening performed utilizing TEX cells in response to one and three rounds of 1 Gy γ-IR (Fig. 2A) (*32, 33*). ‘Leukemia stem cell (LSC)-like’ human hematopoietic cell line TEX was generated via TLS-ERG leukemia fusion oncogene expression in cord-blood derived hematopoietic stem and progenitor cells (HSPCs), which maintains functional heterogeneity, cytokine dependency, and a functional p53 pathway (*34, 35*). Interestingly, *KDM6A* knockdown, similar to loss of other crucial DNA repair genes, significantly impaired proliferation in two out of three clones, suggesting a radioprotective function (Figs. 2A-B). Treatment of U937 cells using γ-IR induced HR gene expression (Fig. 2C and fig. S2E). In contrary, induction of DDR gene and protein expression was significantly impaired in KDM6A or KDM6B deficient AML cell lines, which was accompanied with an altered cell survival and proliferation (Fig. 2C and figs. S2E-H). Ectopic expression of full length KDM6A, but not JmjC mutant, restored DDR gene expression (fig. S2I). Earlier we demonstrated that treatment of AML cells using a KDM6 small molecule inhibitor GSK-J4 causes a selective increase in H3K27me3 (*4*). KDM6A primarily plays tumor suppressor role in demethylase independent mechanisms (*9, 14*). However, we and others reported that KDM6 inhibition in AML cells, with intact KDM6 expression, using GSK-J4 attenuates leukemia cell survival and leukemia development (*4, 12*). Apart from KG1a, all cells displayed IC_50_ greater than 2 µM, a dose we used as a sub-lethal concentration for subsequent experiments (figs. S3A-B). Similar to KDM6 deficiency, treatment with GSK-J4 abrogated the expression of HR and NHEJ genes, highlighting KDM6 demethylase-dependent function in DDR gene regulation (Figs. 2D-E and figs. S3C-J). There was a substantially higher NHEJ rate compared to HR, and KDM6 inhibition compromised HR activity more than NHEJ (figs. S4A-B). KDM6A deficient AML cells, regardless of *TP53* mutation status, revealed an elevated double-stranded DNA break with an attenuated γH2A.X in response to genotoxic stress (Figs. 2F-G). KDM6 inhibition in U937 cells revealed a slightly elevated basal γH2A.X (fig. S4C). In agreement, radiation exposure induced a time-dependent increase in γH2A.X and p-ATM in vehicle treated AML cells, whereas KDM6 blockade showed an impaired γH2A.X and p-ATM induction (fig. S4C). In addition, there was significant transcriptional downregulation of all three MRN components, Mre11a, Rad50 and Nbn along with loss of DSB transducer Atr in *Utx^-/-^* AML cells (fig. S4D). Consistent with these findings, KDM6 knockdown in human AML also caused a reduced expression of MRN (fig. S4E). Collectively, these results underscore that KDM6 proteins play a critical role in maintaining an elevated expression of DSB recognition genes in AML cells, and KDM6 deficiency or inhibition causes an impaired DSB repair response.

**Fig. 2.**
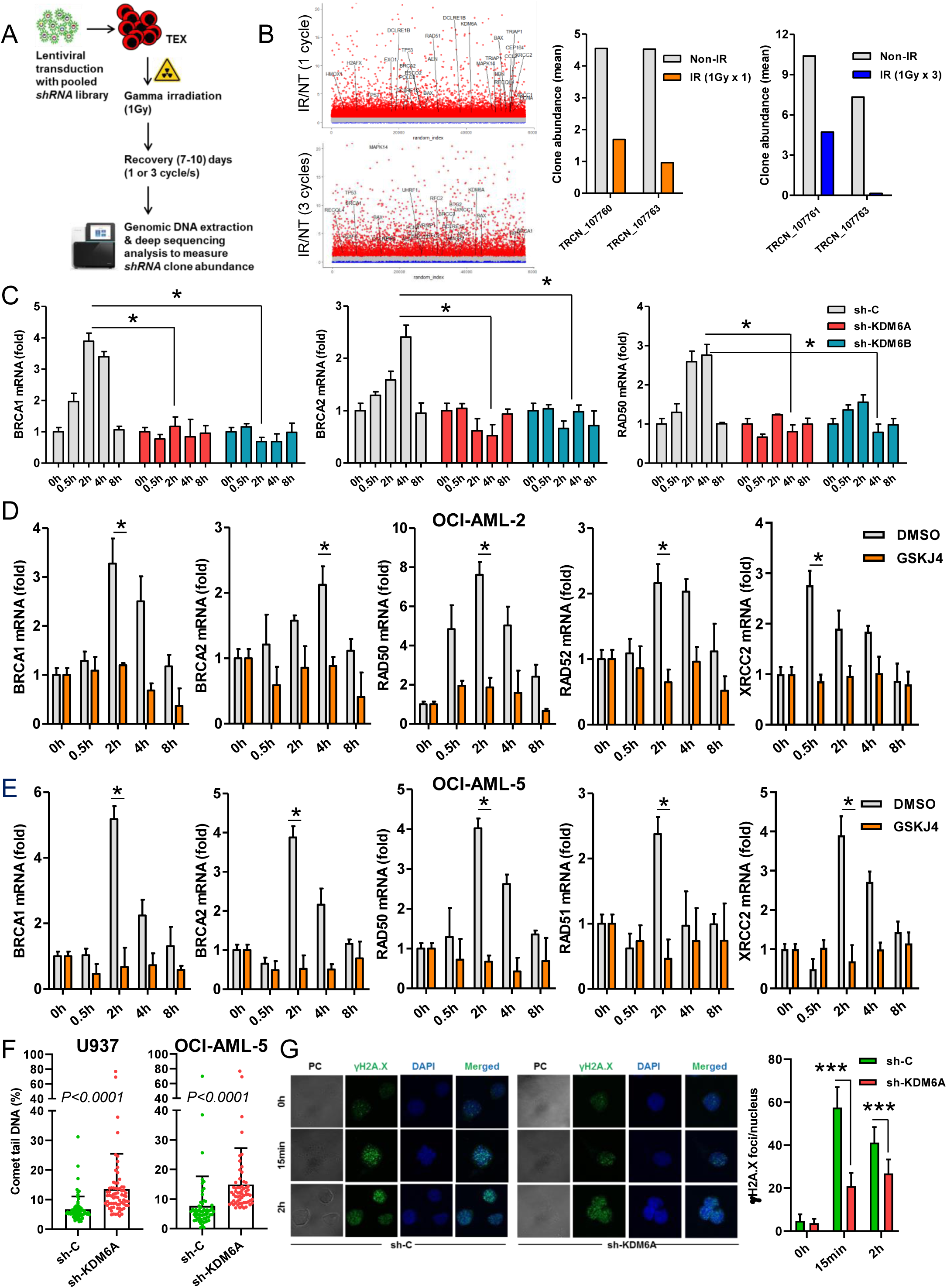
Loss of KDM6 in AML cells impairs DDR gene expression and double stranded break (DSB) repair. (A) Schema representing screening assay for radiosensitive genes using pooled targeted lentiviral *shRNA* library in TEX leukemia cells. (B) Scatter plots showing distribution of KDM6A along with a few other known DNA repair associated genes with respect to the overall gene sets analyzed in the *shRNA* screening in response to 1 round *(upper panel)* or 3 rounds *(lower panel)* of radiation-recovery cycles. The values on the Y-axes denote the ratio [IR (treatment)/NT (control)] of individual *shRNA* corresponding to each gene. Analysis of clone abundance (average of four replicates) of KDM6A targeting *shRNA* clones after 1 round or 3 rounds of γ-IR (1 Gy) and recovery cycles *(right two panels)*. (C) qRT-PCR analysis (normalized to 0h) of HR genes in control and KDM6 deficient U937 cells treated with 10 Gy of γ-IR (n=2). (D) qRT-PCR analysis (normalized to 0h) of HR genes in DMSO (control) and GSK-J4 treated OCI-AML-2 cells treated with 10 Gy γ-IR (n=2). (E) qRT-PCR analysis (normalized to 0h) of HR genes in DMSO and GSK-J4 treated OCI-AML-5 cells irradiated with 10 Gy (n=2). (F) Neutral comet assay showing distance of comet tail measured in control or KDM6A deficient AML cells after 2 hr of treatment with 10 Gy of γ-IR (n=2). (G) Immunofluorescence analysis *(left)* and quantitation *(right)* of γH2A.X foci per nucleus (n=40-50) in control or KDM6A deficient AML cells at different time points after treatment with 10 Gy of γ-IR (n=2). qRT-PCR values were normalized to GAPDH. Data are representative of two to three independent experiments. Statistics were calculated with Student’s t-test; error bars represent means ± SD. \**P* < 0.05 or \*\*\**P* < 0.001 were considered to be statistically significant.

### KDM6A regulates chromatin accessibility and transcriptional activation at DDR loci

To understand the mechanism of KDM6A mediated DDR gene regulation, we conducted qChIP experiments. There was a significant enrichment of KDM6A at the transcription start sites (TSS) and promoter-proximal elements of BRCA and RAD family genes in AML cells (Fig. 3A and fig. S4F). KDM6A downregulation was associated with a concomitant increase in occupancy of H3K27me3 at these loci in both untreated and radiation treated AML cells, further indicating demethylase-dependent transcriptional regulation (Fig. 3B and fig. S4G). γ-IR caused a reduction in H3K27me3 at the HR promoter-proximal loci in control cells, however, a similar decrease in locus specific H3K27me3 occupancy was either absent or negligible in KDM6A deficient cells (Fig. 3B and fig. S4G). KDM6A has been shown to functionally interact with SWI/SNF ATP-dependent chromatin remodeling complex to regulate chromatin accessibility, and influence gene expression (*1, 9, 36*). We had previously conducted ChIP-seq with the SMARCC1 (BAF155) core subunit of SWI/SNF in primary AML samples (GSE108976) (*11, 37*). Reanalysis of genes which showed enrichment for both SMARCC1 and KDM6A revealed a substantial overlap between SMARCC1 and KDM6A targets (2785 genes; dataset S3, *P* < 0.05), with the majority of these co-occupied genes also showing enrichment of H3K27ac, while being devoid of H3K27me3 (1676 genes) (Fig. 3C and dataset S4). These KDM6A targets that include DNA repair genes may represent potential candidates for co-regulation by KDM6A demethylase and SWI/SNF (Fig. 3D). qChIP further demonstrated that concomitant with an increase in KDM6A occupancy there was a significant enrichment of SMARCC1, CBP and H3K27ac at the TSS and promoter regions of BRCA and RAD genes in response to radiation (Fig. 3E). SMARCC1 enrichment along with CBP and H3K27ac was also observed at the promoter region of KDM6A itself (Fig. 3F), suggesting KDM6A and SWI/SNF cooperation in DDR gene regulation.

**Fig. 3.**
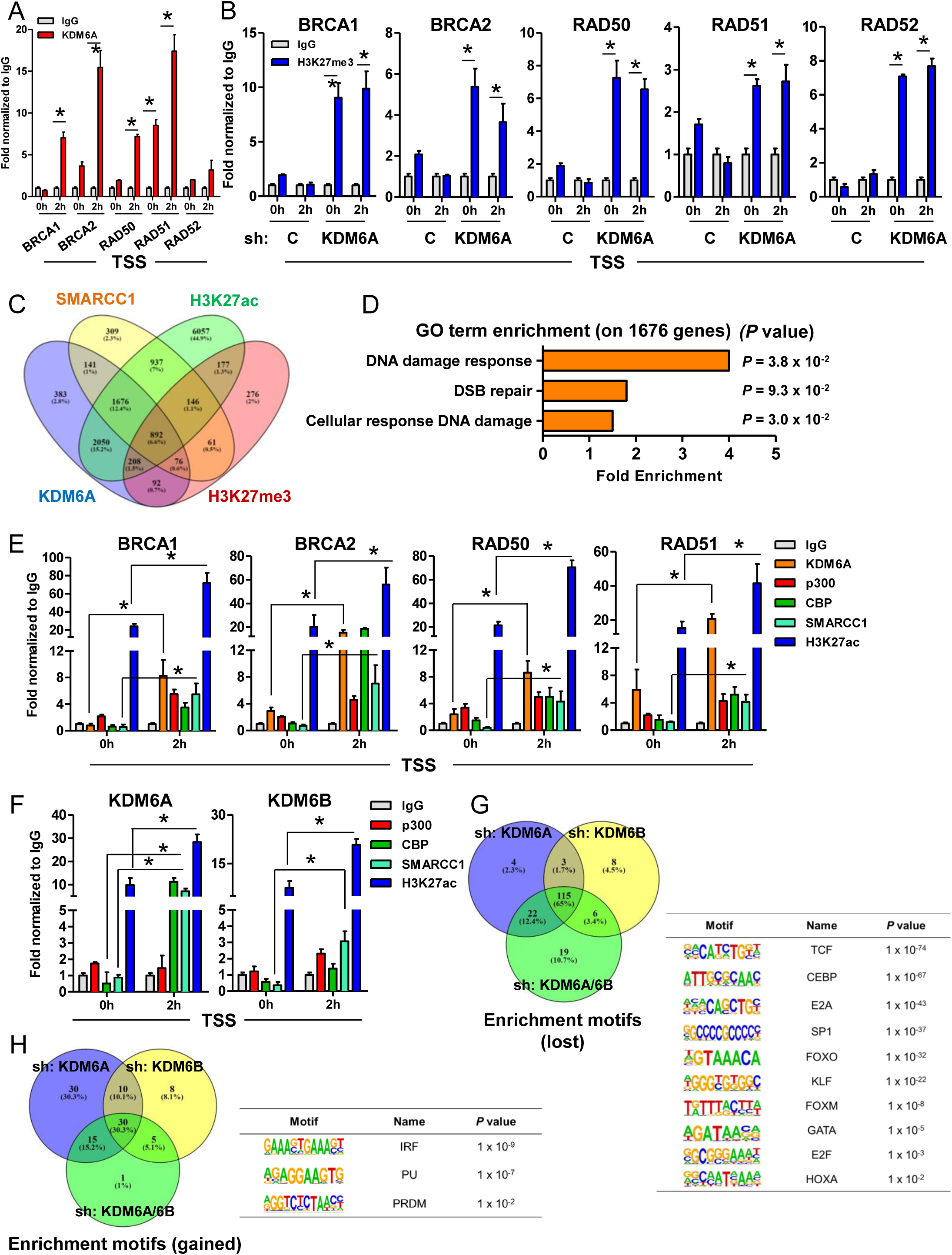
KDM6A regulates chromatin architecture at DDR loci. (A) qChIP analysis showing KDM6A chromatin occupancy on transcription start sites (TSS) of HR genes in U937 cells treated with 10 Gy of γ-IR. (B) qChIP analysis showing H3K27me3 chromatin occupancy on TSS of HR genes in control and KDM6A deficient U937 cells treated with 10 Gy of γ-IR. (C) ChIP-seq venn diagram analysis representing co-occupancy of KDM6A, SMARCC1 and H3K27ac (excluding H3K27me3) in primary AML. (D) GO term analysis of the 1676 co-occupied genes from (C). (E) qChIP analysis showing chromatin occupancy and cooperation of KDM6A and SMARCC1 (BAF155 subunit of the SWI/SNF complex) on TSS of HR genes in U937 cells treated with 10 Gy of γ-IR. (F) qChIP analysis showing chromatin occupancy on TSS of KDM6A and KDM6B in U937 cells treated with 10 Gy of γ-IR. (G) ATAC-seq Motif analysis, of transcription factors associated with DDR gene regulation, representing a significant loss in chromatin accessibility in KDM6A deficient U937 cells. (H) ATAC-seq Motif analysis showing a gain in chromatin accessibility in KDM6A deficient AML. qChIP values were normalized to IgG. Data are representative of two independent experiments. Statistics were calculated with Student’s t-test; error bars represent means ± SD. \**P* < 0.05 was considered to be statistically significant.

To interrogate changes in chromatin accessibility on KDM6 loss, we performed bulk ATAC-seq in KDM6A and KDM6B deficient AML cells and compared them with unaltered control cells. In concordance with transcriptional activation function of KDM6, the number of transcription factor (TF) motifs enriched in control cells that lost accessibility in KDM6 deficient cells was much higher than the number of motifs, which gained accessibility in KDM6A/B deficient AML cells (Figs. 3G-H). Motif comparison revealed greater than 90% overlap in KDM6A or KDM6B deficient cells. There was significant loss in the binding potential of TCF, CEBP, FOXO and HOXA family, which usually promote HR gene expression (Fig. 3G). Alternatively, there was increase in binding potential of IRF, PU and PRDM (Fig. 3H), which have been shown to suppress DDR and induce genomic instability. Collectively, these results indicate that changes in chromatin accessibility correlate with lower abundance of TF binding sites required for optimal DNA gene regulation. Together this may account for the observed repression of DDR gene expression in KDM6 deficient AML, thus compromising DNA repair.

### KDM6A loss renders AML cells sensitive to PARP inhibition

We next assessed whether reduced KDM6 levels would sensitize AML cells to inhibition of PARP-1 signaling. Analysis of OHSU AML (n=672), containing *de novo* and relapsed AML cases with varying molecular subtypes, showed a significant inverse correlation between KDM6A and PARP-1 expression (Figs. 4A-B and fig. S4H). There was no major change in PARP1 expression in KDM6 deficient U937 cells (fig. S4I). PARP inhibition using olaparib for 72 hr decreased intracellular PAR level and induced apoptosis in AML cells (figs. S5A-B). In general treatment with olaparib at concentrations below IC_50_ caused a cytostatic rather than cytotoxic effect; significant cell apoptosis was observed only at higher concentrations (fig. S5B). Drug dose response analysis indicated that AML cells co-treated with GSK-J4 were significantly more sensitive to olaparib compared to the controls (Fig. 4C and fig. S5C). Except KG1a cells, both *TP53* wild type (OCI-AML-2, OCI-AML-5, MOLM-13) and *TP53* mutated (U937, NB4) AMLs were susceptible to olaparib in response to KDM6 inhibition (Fig. 4C). Similarly, deficiency of KDM6A sensitized AML cells to PARP inhibition (Fig. 4D). Additionally, KDM6 loss caused a differential sensitivity of select AML subtypes to a conventional chemotherapeutic agent like AraC, although daunorubicin treatment did not appreciably alter AML sensitivity (figs. S5D-F).

**Fig. 4.**
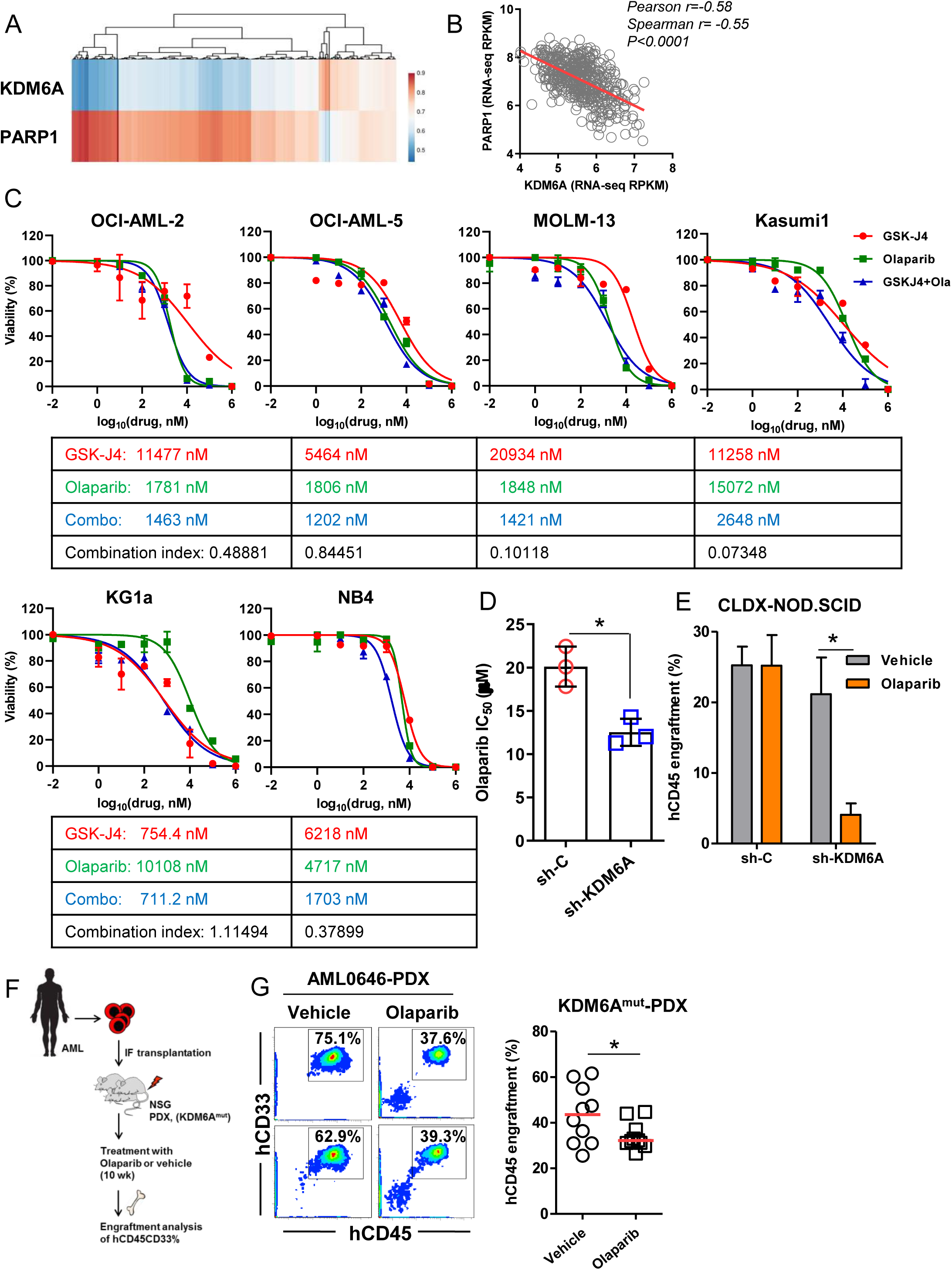
KDM6A deficiency sensitizes AML to PARP inhibition. (A) KDM6A and PARP1 mRNA expression z-scores (RNASeq v2 RSEM) heatmap cluster from OHSU AML dataset. (B) Gene expression correlation analysis of PARP1 with KDM6A in OHSU AML cohort (n=672). (C) Percent viability of AML cells treated with varying doses (from 1 nM to 1 mM) of GSK-J4 alone *(red)* or olaparib alone *(green)* or in combination *(blue)* for 72 hr. Data represent average of two to three independent experiments with similar results. IC_50_ values are tabulated and combination index (Ci) at ED_50_ was calculated using CompuSyn v 1.0. Ci < 1 was considered as drug synergism. (D) IC_50_ of olaparib of control or KDM6A deficient AML cells cultured for 48 hr. Data represent average of two to three independent experiments with similar results. (E) Bone marrow engraftment analysis of human CD45^+^ cells in NOD.SCIDs after treatment with vehicle or olaparib (n=5 for each treatment group). (F) Schema representing bone marrow engraftment analysis performed in *KDM6A* mutant AML patient-derived xenografts (PDX) in response to PARP inhibition. (G) Flow cytometry contour plots *(left)* and quantitative analysis *(right)* showing engraftment of human CD33^+^CD45^+^ cells in NSG mice after being treated with vehicle or olaparib (n=5 for each treatment group). Statistics were calculated with Student’s t-test; error bars represent means ± SD. \**P* < 0.05 was considered to be statistically significant.

Analysis of the Beat AML dataset indicated that cells with lower KDM6A expression may harbor *FLT3-ITD* mutation (fig. S6A). In agreement we observed that *FLT3-ITD* expressing KDM6A deficient AML cells were relatively more sensitive to olaparib compared to the controls (fig. S6B). To investigate olaparib sensitivity *in vivo* we transplanted control and KDM6A deficient U937 into NOD.Cg-*Prkdc^scid^*/J (NOD.SCID) mice (fig. S6C). KDM6A loss alone did not affect the overall engraftment potential. In support of our prediction, compared to vehicle treated cells, olaparib administration resulted in a significant decrease in the engraftment of KDM6A deficient, but not control, AML (Fig. 4E). To further confirm, we established AML patient-derived xenograft models carrying KDM6A nonsense mutation implicated in relapse (Fig. 4F). There was a significant reduction of human CD45^+^CD33^+^ cells in the bone marrow in mice treated with olaparib compared to vehicle treated group (Fig. 4G). Together these results suggest that KDM6A loss increases sensitivity of AML cells to PARP inhibition.

### Deficiency of KDM6A increases susceptibility of AML to BCL2 blockade

BCL2 inhibitor venetoclax has shown promise in the clinical setting, although a majority of the initial responders relapse (*38, 39*). Beat AML analysis indicated that monocytic (Mono) AML cases associate with venetoclax resistance (*40*), as well KDM6A^hi^ expressing male AMLs are relatively more tolerant to venetoclax (Figs. 5A-B). We re-analyzed the available RNA-seq dataset from venetoclax resistant Mono-AML ROS^low^ LSCs, and compared with venetoclax sensitive Prim-AML Ros^low^ LSCs (*41*). In agreement with earlier findings Mono-AML showed a relatively lower BCL2, and there was a significant increase in BCL2A1 expression in Mono-AML compared to Prim-AML (Fig. 5C and fig. S6D). BCL2A1 is a predictive biomarker of venetoclax resistance in AML and induces resistance to BCL2 inhibitor ABT-737 in CLL (*42, 43*). Consistent with these findings, the OHSU AML dataset further suggested a positive correlation between KDM6A and BCL2A1 expression (Fig. 5D). KDM6A downregulation induced BCL2, which was accompanied with a concomitant decrease in BCL2A1 gene expression (Figs. 5E-F and figs. S6E-G). We further reanalyzed available transcriptome dataset (FDR: 0.05; Log_2_FC: > 1.25) (*14*) of *KDM6A*-null THP1 cells ectopically expressing full length *KDM6A* or various domain mutants of *KDM6A* (fig. S6H). Gain in full-length *KDM6A*, but not TPR or JmjC deletion mutants to some extent, induced BCL2A1 expression in THP1 cells (fig. S6I). Although cIDR-deleted *KDM6A* mutant did not restraint BCL2A1 induction, chimeric IDRs partly restored BCL2A1 expression (fig. S6I). In addition, qChIP analysis performed in our AML cell lines panel identified KDM6A occupancy at BCL2 or BCL2A1 TSS and promoter regions (figs. S6J-K). While KDM6A deficiency resulted in increased occupancy of p300 and H3K27ac at BCL2 promoter, there was increased H3K27me3 and reduced p300 at BCL2A1 loci in KDM6A deficient AML cells (Fig. 5G and fig. S6L). Additionally, corroborating these results GSK-J4 treatment at respective IC_50_ doses induced BCL2 expression in select AML subtypes (except MOLM-13 and Kasumi1 cells) (Fig. 5H and figs. S6M-N). Collectively, these findings indicate that KDM6A differentially regulates BCL2 family gene expression, and KDM6A loss correlates with BCL2 induction.

**Fig. 5.**
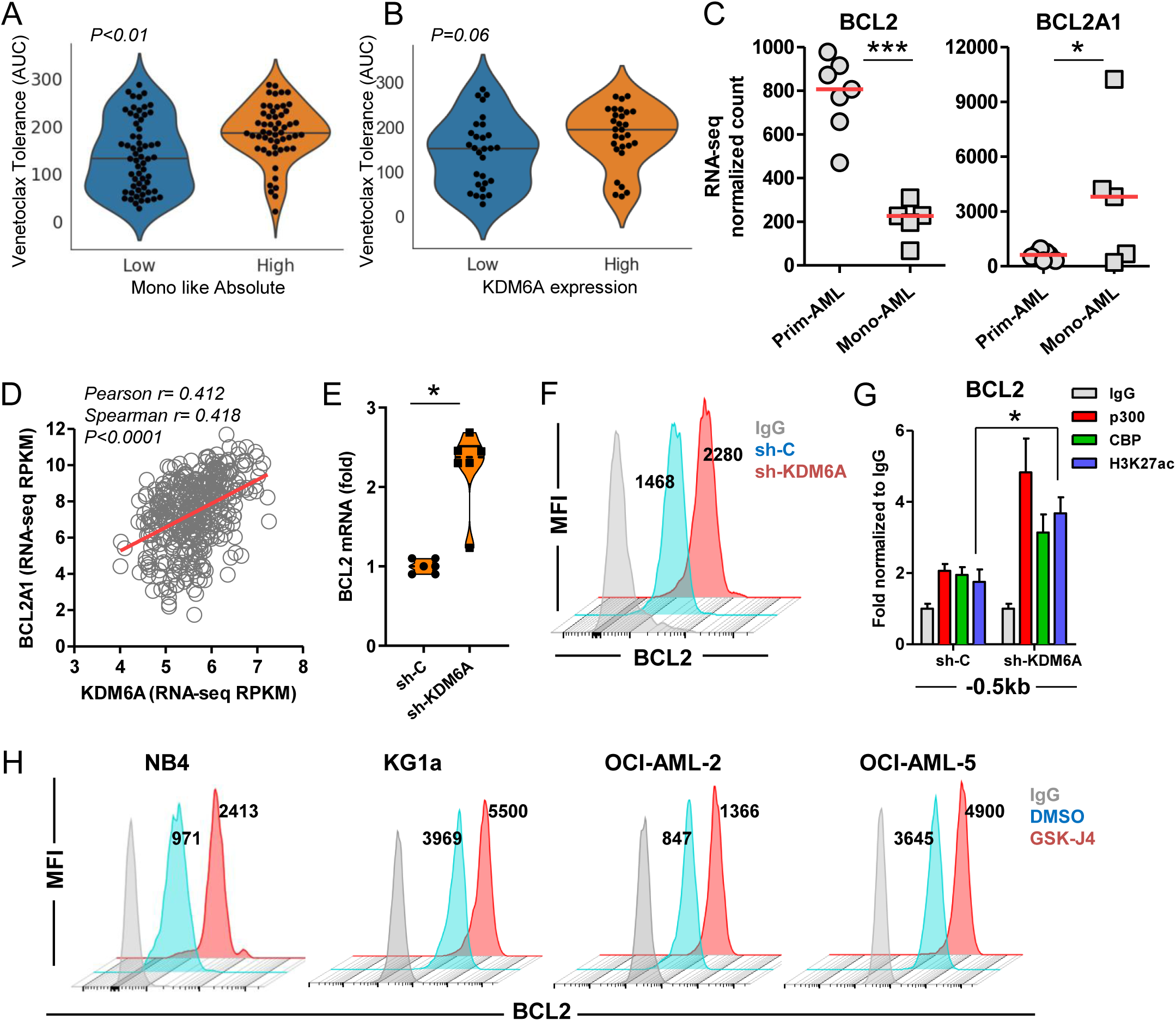
KDM6A associates with BCL2 and BCL2A1 expression. (A) Venetoclax tolerance (AUC) based on abundance of monocytic (Mono-) AML from Beat AML (n=702). (B) Venetoclax tolerance analysis performed between KDM6A-low and KDM6A-high expressing male AML from Beat AML cohort. (C) RNA-seq analysis showing expression of BCL2 and BCL2A1 between primitive (Prim-) AML (n=7) and monocytic (Mono-) AML (n=5) ROS^low^ LSCs. (D) Gene expression correlation analysis between KDM6A and BCL2A1 in OHSU AML dataset (n=672). (E) qRT-PCR analysis of control or KDM6A deficient U937 cells. Error bars represent means ± SEM. (F) Flow cytometry staggered histogram plots showing BCL2 expression in control or KDM6A deficient U937 cells. (G) qChIP analysis showing chromatin occupancy at BCL2 promoter region (−0.5 Kb) in control and KDM6A deficient U937. (H) Flow cytometry staggered histogram plots showing BCL2 expression in AML cells treated with DMSO *(blue)* or GSK-J4 *(red)* at respective IC_50_ concentrations for 48 hr. qRT-PCR and qChIP values were normalized to GAPDH and IgG, respectively. Data represent two to three independent experiments. Statistics were calculated with Student’s t-test; error bars represent means ± SD if not specified otherwise. \**P* < 0.05 or \*\*\**P* < 0.001 were considered to be statistically significant.

BCL2 induction commonly associates with venetoclax function (*44*). GSK-J4 mediated BCL2 induction in AML subtypes further prompted us to interrogate venetoclax sensitivity. Indeed, dose response analysis revealed that *TP53* wild type (OCI-AML-2, OCI-AML-5) as well as *TP53* mutant (NB4, KG1a) AML cells co-treated with either varying doses of GSK-J4 or constant doses of GSK-J4, set at half of the IC_50_ concentrations of respective cell types, were significantly more sensitive to venetoclax compared to the monotherapies alone (Fig. 6A). Although MOLM-13 partially responded to this combination, Kasumi1 cells did not show any effect (Fig. 6A). Similarly, deficiency of KDM6A also sensitized AML cells to BCL2 inhibition (Fig. 6B). In addition, olaparib treatment resulted in an increase in mitochondrial membrane potential (MMP) in KDM6A deficient AML cells compared to control cells (Fig. 6C). Inhibition of KDM6 and PARP also resulted in an increase in MMP in AML cells (Fig. 6D). Furthermore, MV4-11 venetoclax resistant (Ven-res) cells showed a decrease in MMP and ROS level compared to venetoclax sensitive (Ven-sen) group (figs. S7A-D). KDM6 inhibition restored ROS in MV4-11 Ven-res cells, which was further increased in presence of olaparib (figs. S7C-D). Intriguingly, we provide evidence that changes in BCL2 expression and mitochondrial activity associated with KDM6A loss, may account for venetoclax tolerance in AML.

**Fig. 6.**
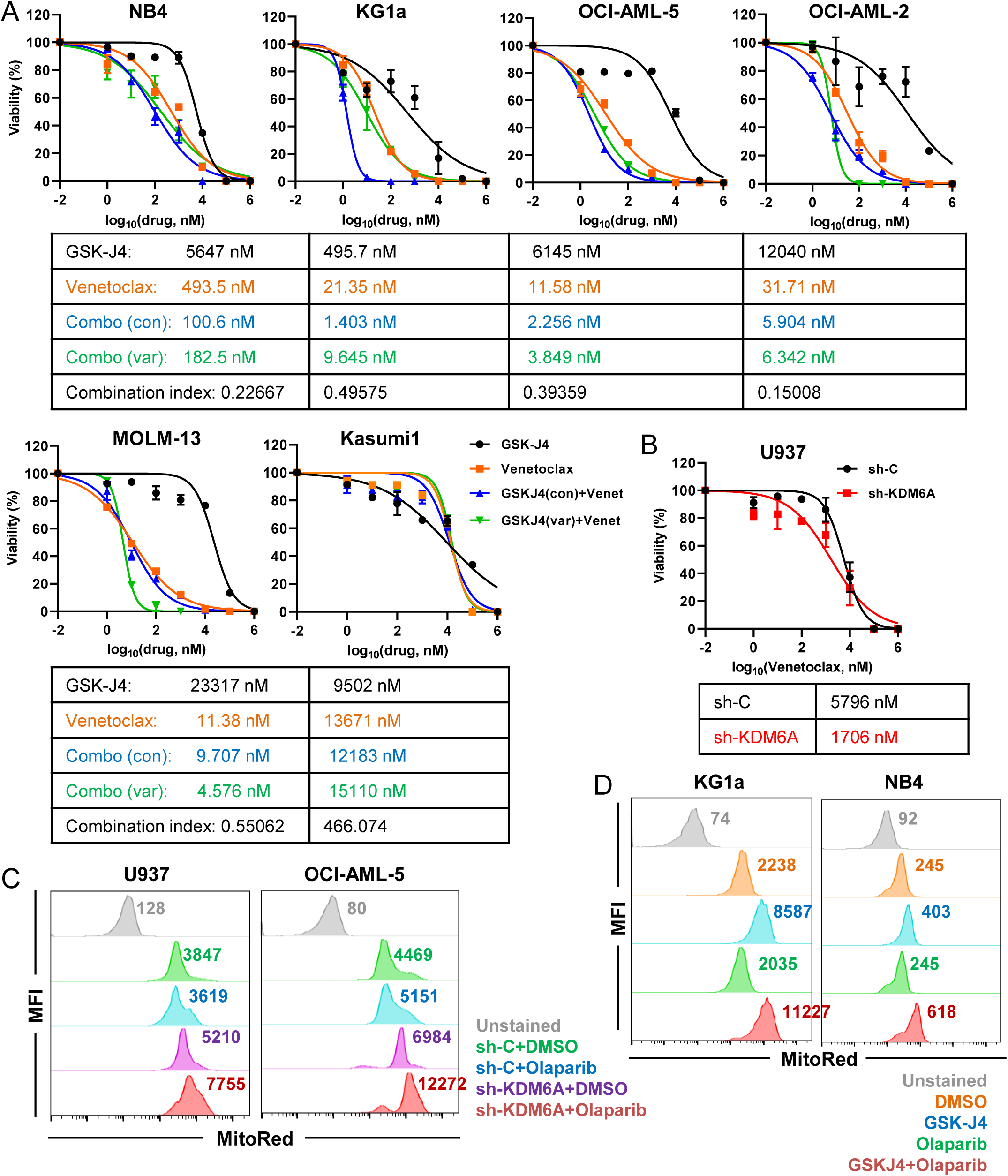
Attenuation of KDM6 increases AML susceptibility to BCL2 blockade. (A) Percent viability of AML cells treated with varying doses (from 1 nM to 1 mM) of GSK-J4 alone *(var, black)* or venetoclax alone *(orange)* or in combination *(green)* for 72 hr. Additional combination *(blue)* line represents a constant dose of GSK-J4 *(con)* used at half of the respective IC_50_ concentrations. Data represent average of two to three independent experiments with similar results. IC_50_ values are tabulated and Ci at ED_50_ was calculated using CompuSyn v 1.0. Ci < 1 was considered as drug synergism. (B) Percent viability of control or KDM6A deficient U937 cells treated with varying doses (from 1 nM to 1 mM) of venetoclax for 48 hr. Data represent average of two to three independent experiments with similar results. (C) Flow cytometry histogram overlay analysis showing mitochondrial membrane potential (MMP) in control or KDM6A deficient AML cells treated with olaparib (10 µM) or DMSO for 48 hr. (D) Flow cytometry histogram overlay analysis showing MMP in AML cells treated with DMSO *(orange)* or GSK-J4 *(blue)* or olaparib *(green)* or a combination of GSK-J4 and olaparib *(red)* at respective IC_50_ doses for 72 hr.

### Dual inhibition of PARP and BCL2 synergizes in AML

Next we investigated whether inhibition of PARP and BCL2 would have a combination effect in controlling AML cell survival. Co-treatment of olaparib and venetoclax were superior in inhibiting AML cell viability compared to the monotherapies alone (Fig. 7A). Combination of olaparib and venetoclax showed synergistic effects in reducing cell survival in select AML subtypes including OCI-AML-2, OCI-AML-5, KG1a, NB4 and U937 (Fig. 7A). We did not observe drug synergism in MOLM-13, Kasumi1 and HL60 cells (Fig. 7A). Similarly, dual inhibition of PARP and BCL2 signaling induced apoptosis in AML (Fig. 7B and figs. S7E-F). Interestingly, KDM6A deficient AML cell lines were even more sensitive to the combination therapy (Figs. 7C, D).

**Fig. 7.**
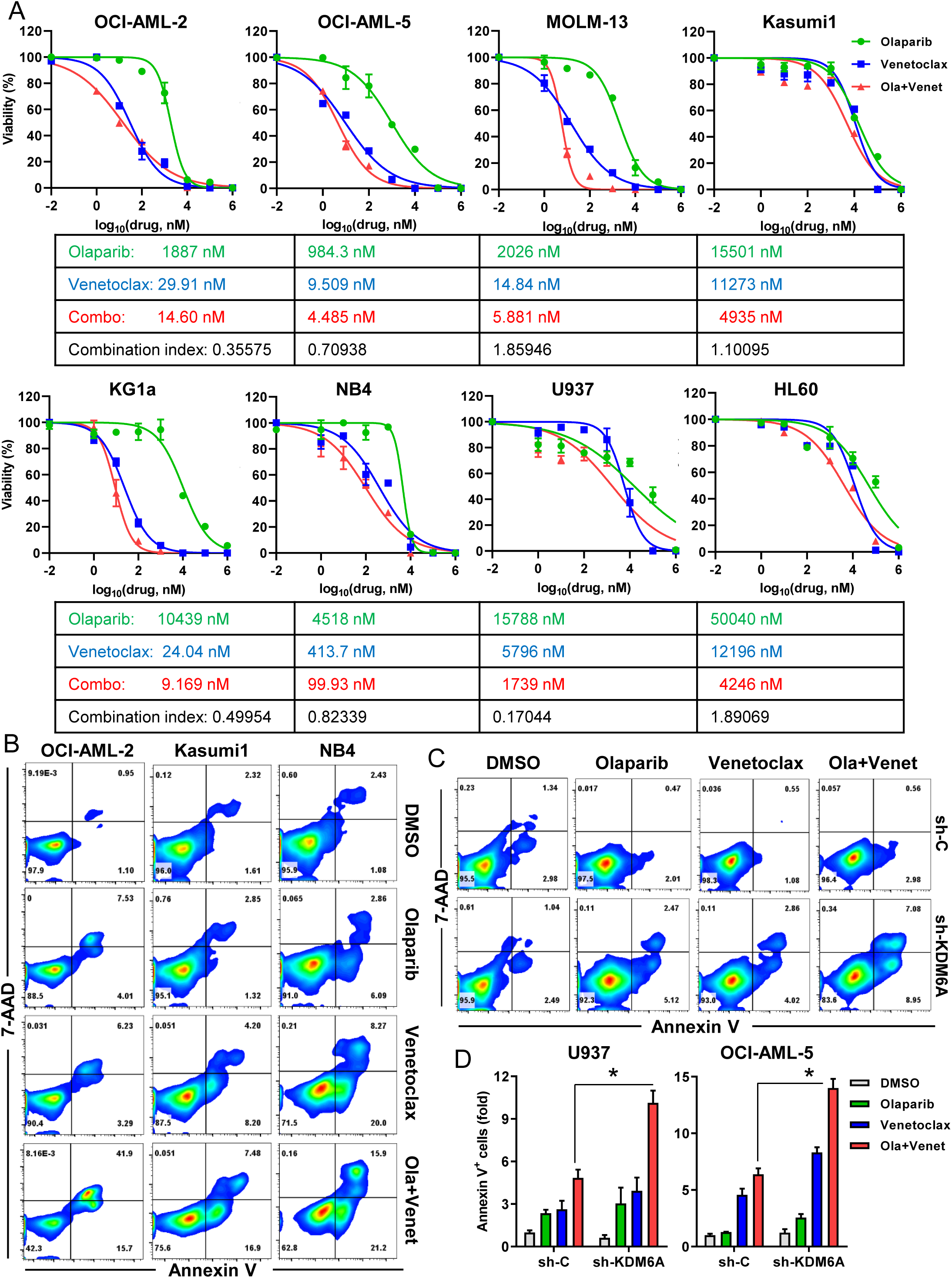
PARP inhibition synergizes with BCL2 blockade in AML. (A) Percent viability of AML cells treated with varying doses (from 1 nM to 1 mM) of olaparib alone *(green)* or venetoclax alone *(blue)* or in combination *(red)* for 48 hr. Data represent average of two to three independent experiments with similar results. IC_50_ values are tabulated and Ci at ED_50_ was calculated using CompuSyn v 1.0. Ci < 1 was considered as drug synergism. (B) Flow cytometry analysis showing apoptosis in AML cells treated with either olaparib or venetoclax or a combination of olaparib and venetoclax at respective IC_50_ concentrations for 48 hr. (C) Flow cytometry analysis showing apoptosis of control and KDM6A deficient U937 treated with either DMSO, olaparib, venetoclax, or a combination of olaparib and venetoclax at IC_50_ doses for 48 hr. (D) Fold change in apoptosis in control or KDM6A deficient AML cells treated with either olaparib or venetoclax or combination for 48 hr.

To further confirm, we argued that KDM6A silencing may not necessarily mimic pathologically occurring *KDM6A* mutations. Therefore, we compared drug sensitivities in primary AML samples carrying either wild type *KDM6A* or different acquired domain mutants of *KDM6A* (Fig. 8A). Corroborating our findings, olaparib and venetoclax treatment showed a stronger synergistic effect in inhibiting viability of *KDM6A*-domain mutant primary AML samples compared to *KDM6A*-wild type (WT) cases (Figs. 8B-C). Although we could not test drug efficacy in cIDR-mutant, both TPR and JmjC mutants had dramatic loss of cell viability in response to olaparib and venetoclax (Fig. 8C). *NPM1^mut^* AML848978 only showed a marginal response to the combination (Fig. 8C). Similarly, combination of PARP and BCL2 inhibition led to an increase in apoptosis in *KDM6A* mutant primary AML cells compared to the wild type control cells (Fig. 8D and fig. S7G). Normal HSPCs were relatively more tolerant to olaparib (average IC_50_: 13.56 µM compared to 0.42 µM in *KDM6A^mut^* primary AML) and venetoclax (average IC_50_: 11.46 µM compared to 0.18 µM in *KDM6A^mut^*primary AML) (figs. S7H-I). Overall, KDM6A loss had the most profound effect by compromising DNA damage response and inducing BCL2, thus rendering AML cells sensitive to PARP and BCL2 blockade (Fig. 8E). In sum, we provide evidence and rationale supporting pre/clinical testing of the novel combination targeted therapy for human AML, and posit KDM6A as an important regulator in determining therapeutic efficacy in AML subtypes.

**Fig. 8.**
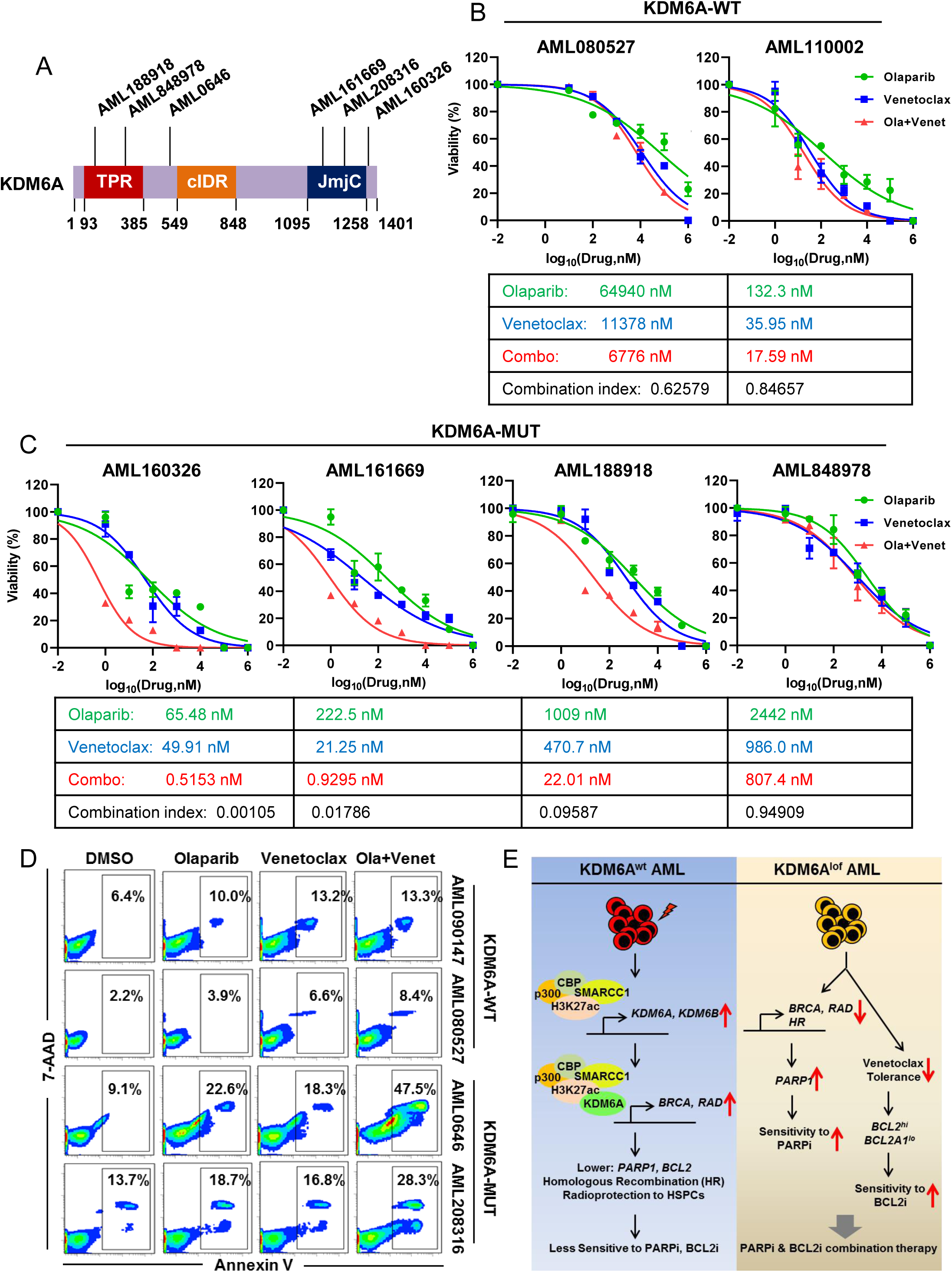
KDM6A-domain mutant primary AML cells are even more sensitive to combination of PARP and BCL2 blockade. (A) Schema showing primary AML cells carrying different *KDM6A*-domain mutants used in our study. (B-C) Percent viability of primary AML cells, carrying (B) wild type or (C) mutant *KDM6A*, treated with varying doses (from 1 nM to 1 mM) of olaparib alone *(green)* or venetoclax alone *(blue)* or in combination *(red)* for 48 hr. Data represent average of two independent experiments with similar results. IC_50_ values are tabulated and Ci at ED_50_ was calculated using CompuSyn v 1.0. Ci < 1 was considered as drug synergism. (D) Flow cytometry contour analysis showing apoptotic cell populations in *KDM6A*-wild type *(upper panels)* and *KDM6A*-mutant *(lower panels)* primary AML patient-derived mononuclear cells in response to either DMSO, olaparib, venetoclax, or a combination of olaparib and venetoclax for 48 hr. (E) Overall schema represents KDM6 deficiency induced sensitization of PARP and BCL2 blockade in AML

## DISCUSSION

In this study we illustrate a mechanistic connection between KDM6 function to impaired DNA repair and BCL2 dependence in AML cell survival. Although venetoclax tolerance is primarily determined by BCL2 expression, and BCLA1 associates with resistance, molecular epi/genetic regulation of these two key proteins is unknown. We provide the first evidence in support for a central regulation integrated by KDM6A demethylase towards BCL2 and BCL2A1 expression important for AML pathogenesis. Our findings that KDM6A was an important regulator for determining efficacy of both PARP and BCL2 blockade; provide support for molecular subtype guided combination targeted therapy for AML. Venetoclax in combination with other small molecule inhibitors has shown better efficacy than venetoclax alone (*38, 45*). Combination therapy using venetoclax with Complex I inhibitor, MAPK pathway inhibitor or cytarabine has shown promise in pre-clinical AML models (*46–48*). In addition, combining KDM6 pharmacological inhibition with venetoclax has been shown to be effective in MYCN amplified neuroblastoma (*49*). Although BCL2 inhibition has been used in combination with hypomethylating agents, their effectiveness in synergy with PARP blockade in AML remains unexplored. In agreement with our findings an ongoing study indicates that PARP Inhibition using talazoparib can enhance anti-leukemic activity of venetoclax in preclinical human AML models [Blood (2021) 138 (Supplement 1): 1176]. Therefore, stratifying AML patients based on KDM6A mutation or expression analysis, should aid in improving therapeutic combinations.

While HR mediated DSB repair is indispensable for survival of MLL-AF9 transformed AML, most therapy-related AML have an abnormal DSB response (*26, 50*). KDM6 inhibition was shown to induce DNA damage in differentiating ES cells (*51*). Inhibition of KDM6 catalytic activity impairs HR mediated DSB repair and augments radiosensitivity in solid tumors (*31, 52*). Therefore, unlike the demethylase-independent, tumor suppressor function of Utx in AML development, DDR gene regulation is dependent on KDM6A demethylase function (*9, 11*). In addition, we provide evidence for KDM6A and SWI/SNF cooperation in regulating DDR gene expression. Different subunits of the SWI/SNF complex have been implicated to have non-transcriptional roles in DSB repair. For example, the *BRG* bromodomain was shown to directly interact with γ-H2A.X and promote chromatin remodeling around DSBs (*53*). Also *ARID2* facilitates *RAD51* recruitment and HR-mediated repair (*54*).

Tumors deficient in BRCA genes have suppressed repair system and respond to PARP inhibition (*27*). However, AML patients have a low mutational burden for BRCA, and only select subtypes have been shown to have defective DDR that respond well to PARP inhibition. AML1-ETO and PML-RARα driven AML have suppressed expression of key HR associated genes, and are sensitive to olaparib, whereas MLL-AF9 harboring AML is HR proficient and insensitive to PARP inhibition (*28*). Only when used in combination with cytotoxic drugs like cytarabine or daunorubicin does MLL-AF9 AML respond to PARP inhibition (*55, 56*). Therefore, inducing a ‘BRCAness’ phenotype, through epigenetic modulation expands the range of AML patients, previously unresponsive to treatment, that might respond to PARP inhibitors. In accordance with this we illustrate that KDM6 attenuation in general sensitizes AML to PARP inhibition.

We also demonstrate altered chromatin accessibility in KDM6 deficient AML. The majority of these changes entailed loss in accessibility to transcription factors (TFs), like TCF and FOXM, supporting KDM6A’s function as a transcriptional activator. Loss of TCF was reported to attenuate DSB repair and sensitizes colorectal cancer cells to radiotherapy (*57*). TCF target NEIL1, a base excision repair gene, is downregulated in KDM6A deficient cells. In addition, FOXM regulates transcription of BRIP1, which cooperates with BRCA1 to promote HR repair (*58*). BRIP1 expression is also downregulated in KDM6A deficient AML. HOXA9 is the mediator of resistance of MLL-AF9 leukemia to olaparib (*28*). It promotes transcription of key HR genes involved in DSB repair, like *MCM9*, *NABP*, *BLM*, *ATM*, *RAD51C*, *RPA1*, *BRCA1* and *BRCA2*. Importantly, among genes downregulated in KDM6A deficient cells are *NABP*, *ATM*, *BRCA1* and *BRCA2*. Additionally, our findings indicate a putative association of olaparib sensitivity with KDM6A expression and *FLT3-ITD* mutation. *FLT3-ITD* AML occurs in about 30% of all AML patients, have a high leukemic burden, poor prognosis and routinely relapse (*59*). FLT3-ITD has been shown to drive increased ROS production, resulting in extensive DNA damage accumulation (*60*). Therefore, together with low levels of KDM6A and impaired HR, it represents a suitable target for PARP inhibition. Indeed it has been demonstrated that *FLT3-ITD* AML is highly sensitive to olaparib (*61*).

Loss of KDM6A expression and acquired resistance for conventional chemotherapy (*8*) led to the impetus to further interrogate potential synthetic lethal vulnerabilities in AML In sum, we present a molecular framework highlighting that absence of KDM6A is an important mediator of compromised DDR in different AML subtypes and determining response to PARP inhibition. Collectively, our results are in agreement with previous findings showing KDM6A tumor suppressor properties. Importantly, our findings greatly extend this field both mechanistically but also in terms of clinical relevance as it not only illustrates efficacy of PARP blockade in KDM6A deficient AML, but it also highlights *proof of concept* for epigenetic modulation guided combination targeted therapy (PARP and BCL2 blockade) in a different subtype of AML where KDM6A expression is upregulated or intact. Although bi-allelic *Utx* deficiency causes evolution to myeloid neoplasms, perhaps minimal KDM6 activity is important for survival of human AML cells similar to what observed in TET2 deficient AML (*62*). Transcriptional adaptation in response to genetic, epigenetic or metabolic perturbations remains a cardinal phenomenon of AML evolution (*63*). Adaptive chromatin remodelling mediated by KDM6 proteins were found to be important for persistence and drug tolerance of glioblastoma stem cells (*64*). Future studies should investigate to what extent KDM6 proteins cooperate with clonal hematopoiesis associated mutational burden and impinge on chromatin topology and epigenomic landscape in AML pathophysiology. KDM6 demethylases have been implicated in solid tumors, and both PARP and BCL2 inhibitors are already being tested in cancer patients, suggesting a broader scope of application. To conclude, KDM6A emerges to be a common regulator for susceptibility of AML to both PARP and BCL2 inhibition, expanding the possibility to characterize effective combination targeted therapy for AML subtypes in pre/clinical settings.

## MATERIALS AND METHODS

### AML patient samples

Peripheral blood specimens were obtained at diagnosis from nine adult AML patients (KDM6A wild type: AML090147, AML080527, AML110002 and KDM6A mutant: AML0646, AML208316, AML160326, AML188918, AML161669, AML848978) according to pre-established guidelines approved by the Research Ethics Board of the Princess Margaret Cancer Centre, University Health Network (UHN), Toronto, Canada and were conducted in accordance with recognized ethical guidelines. All samples were frozen viably and stored long term at -150**°**C. Samples were selected retrospectively based on genotypes, and the details are available in Table S1. Bone marrow (BM) aspirates were also collected from human primary elderly AML patients at Park Clinic, Kolkata, and umbilical cord blood samples were obtained from termed pregnancies from NRS Medical College & Hospital, Kolkata after informed consent and in accordance with guidelines set by Institutional Human Ethics Committee as well as CSIR-Indian Institute of Chemical Biology Institutional Review Board. Sample collection was a part of the routine diagnosis (*4, 11, 37*). Based on histopathology, karyotype and immunophenotyping analysis of the bone marrow aspirates the samples were included in the study. Low-density (1.077g/cc) mono-nuclear cells were obtained from AML BM using Ficoll density gradient centrifugation (Stem cell technologies).

### Cells

OCI-AML-2, OCI-AML-3, OCI-AML-5, OCI-AML-8227, MOLM-13, KG1a, NB4, Kasumi-1 and 5637 cells were generous gifts from Mark D. Minden and John E. Dick (Princess Margaret Cancer Centre, UHN, Toronto, Canada). OCI-AML-21 and venetoclax sensitive (Ven-sen) and resistant (Ven-res) MV4-11 cells were provided by Steven M. Chan (Princess Margaret Cancer Centre, UHN, Toronto). Briefly, Ven-res MV4-11 cells were generated by slowly escalating drug concentration (up to 2000 nM) over a period of 2 months followed by clonal selection (*65*). The resistant MV4-11 clones expressed higher amounts of MCL-1 and BCL2–like protein 1 (BCL2L1, also known as BCL-XL), respectively, compared with their sensitive parental cells. Resistant AML clones did not have a glycine-to-valine mutation (Gly101Val) in the BCL2 gene. OCI-AML-2, OCI-AML-3, MOLM-13 and KG1a were grown in α-MEM medium supplemented with 10% FBS. OCI-AML-5 culture media was supplemented with 10 ng/mL of recombinant GM-CSF (Peprotech) or 5637 conditioned medium. Kasumi-1 was grown in RPMI supplemented with 20% FBS. OCI-AML-8227, which are arranged in a hierarchy of bulk and stem cells with the LSCs enriched in the CD34^+^CD38^−^ fraction, was cultured in X-VIVO-10 supplemented with 20% BIT, 50 ng/mL SCF, Flt3-L, 25 ng/mL TPO, 10 ng/mL IL-6, IL-3 and G-CSF (*66*). All cells were Mycoplasma-free and validated by STR profiling. U937, HL60, THP1, K562, 293T and Phoenix GP cells were obtained from Jose Cancelas (Cincinnati Children’s Hospital Medical Center) and maintained in RPMI 1640 medium supplemented with 10% FBS. All cells were maintained in presence of 100 U/mL Penicillin, 100 μg/mL Streptomycin and 2 mM L-Glutamine (all from Gibco) at 37°C with 5% CO_2_. Primary AML mononuclear cells were cultured using X-VIVO-10 (Lonza) StemSpan SFEM II (Stem Cell Technologies) supplemented with recombinant 50 ng/mL SCF and FLT3-L, 25 ng/mL TPO and 10 ng/mL IL-3, IL-6 and G-CSF (all from Peprotech or Stem Cell Technologies).

### Mice and xenotransplantation studies

Animal experiments were approved by the Princess Margaret Cancer Centre, UHN, Toronto Animal Care Committee and we confirm that all experiments conform to the relevant regulatory and ethical standards. AML patient-derived xeno-transplantation (PDX) experiments were performed in 8 to 12 week-old female *NOD.Cg-Prkdc^scid^Il2rg^tm1Wjl^/SzJ* (NSG) mice (JAX) that were sublethally irradiated with 225 cGy using a ^137^Cs γ-irradiator 24 hr before intrafemoral transplantation. U937 cell line-derived xenograft (CLDX) studies were conducted in 8 to 12 week male or female *NOD.Cg-Prkdc^scid^/J* (NOD.SCID) mice (JAX) that were sub-lethally irradiated with 225 cGy, 24 hr before transplantation. NOD.SCID mice received systemic administration of anti-CD122 antibody (200 µg i.p.) prior to transplantation. Sample size was chosen to give sufficient power for calling significance with standard statistical tests. Intra-femoral injections were performed as described earlier (*67*). For this, mice were anesthetized with isoflurane and the right knee was secured in a bent position to drill a hole into the RF with a 27 gauge needle. Then, 1 x 10^6^ primary AML cells (for PDX) or 100 x 10^3^ wild type or KDM6A deficient U937 cells (for CLDX) were injected in 30 μL PBS using a 28 gauge ½ cc syringe (Becton Dickinson).

For generating AML-PDX model, prior to intrafemoral transplantation human T-lymphocytes were depleted using CD3 depletion kit (Miltenyi Biotech) from AML0646 mononuclear cells. AML0646 (Female, intermediate cytogenetics, 48,XX,+8,+20[16], ITD negative) carries *KDM6A* nonsense mutation (c.1347G>A; p.W449*), and CD34^+^CD38^-^ LSCs (1/2000) and CD34^+^CD38^+^ (1/26,000) fractions (but not CD34^-^ fractions) were previously characterized to engraft into NSGs in limiting dilution analysis. To further validate in this study both unfractionated as well as CD34^+^ fractions of AML0646 successfully engrafted into NSG recipients. After 6 weeks of transplantation, olaparib (Medkoo Biosciences) (50 mg/kg body weight) or vehicle (DMSO/PEG-400/H_2_O) was administered intraperitoneally twice daily for 5 days a week. After 3 weeks of treatment the mice were euthanized and analyzed for bone marrow engraftment of human CD45^+^CD33^+^ cells. For CLDX, after 72 hr of transplantation mice were started treating intraperitoneally with olaparib 50 mg/kg body weight, twice daily for 5 days per week or vehicle for 4 weeks. After the drug treatments, mice were sacrificed to obtain bone marrow (right and left femur together). Bones were flushed in 1 mL PBS + 2.5% FBS and cells were centrifuged at 350 x g for 10 min. Cells were resuspended in 500 μL of PBS + 2.5% FBS, counted in ammonium chloride (Stem Cell Technologies) using the Vi-CELL XR viability analyzer (Beckman Coulter) and analyzed for human CD45^+^CD33^+^PI^-^ (anti-CD45-APC, anti-CD33-PerCP, anti-CD34-APC.Cy7, anti-CD38-PE) engraftment with 96-well high throughput auto-acquisition mode using FACSCelesta (Becton Dickinson).

### Plasmids

KDM6A targeting human *shRNA* expressing plasmids were purchased from OriGene (TL300596C and TL300596D). KDM6B targeting human *shRNA* plasmid was purchased from Sigma (TRCN0000236677). Scrambled vector *sh-Control (sh-C)* was purchased from OriGene (pGFP-C-sh-Lenti, TR30021). Lentiviral packaging constructs PAX2 (12260) and pMD2.G (12259) were purchased from Addgene. *pCMV-HA-UTX* (24168) encoding human *KDM6A* with HA tag and *pCS2-UTX-F-MT2* (40619) encoding enzyme-dead human *KDM6A* were purchased from Addgene. The HR and NHEJ reporter plasmids were kind gifts from Tomasz Skorski (*61*). *FLT3-ITD* lentiviral construct was generously provided by Mark D. Minden (Princess Margaret Cancer Centre, UHN, Toronto).

### Lentivirus preparation and transduction

293T cells were maintained in DMEM supplemented with 10% FBS, 100 U/mL Penicillin, 100 mg/mL Streptomycin, and 2 mM L-Glutamine (all from Thermo Fisher Scientific) at 37°C with 5% CO_2_. For production of lentiviral particles, cells were transfected with target DNA, PAX2 (Addgene) and pMD2.G (Addgene) plasmids using calcium phosphate transfection method keeping cell density at 70% confluency (*11, 37*). Supernatants containing viral particles were collected and concentrated using ultracentrifugation at 25,000 rpm for 90 min at 4°C using (Sorvall WX Ultra 90; Thermo Fisher Scientific). AML cells were transduced with lentiviral particles (MoI=5) expressing non-targeting *shRNA (sh-C)*, KDM6A-targeting *shRNA (sh-KDM6A)* and co-expressing GFP or KDM6B-targeting *shRNA (sh-KDM6B)* in a U-bottom 96-well non-tissue-culture treated plate in presence of polybrene (8 µg/mL) and incubated overnight at 37°C with 5% CO_2_. Cells were centrifuged, resuspended in fresh media and cultured for another 48 hr at 37°C with 5% CO_2_ before proceeding with downstream applications.

### Flow cytometry sorting and generation of KDM6 deficient AML cells

U937 and OCI-AML-5 cells were transduced with lentiviral particles expressing *sh-C* or *sh-KDM6A* and co-expressing GFP. GFP^+^7-AAD^-^ cells were sorted using MoFlo XDP Cell Sorter (Beckman Coulter) or FACSAria III Cell Sorter (Becton Dickinson). Enrichment of the sorted cells was 90% to 95% on average. *sh-KDM6B* construct did not co-express a fluorescent marker. Therefore, *sh-KDM6B* expressing AML cells were selected with puromycin 2 (µg/mL) for 48 hr and subsequently characterized by immunoblot analysis. Similarly, KDM6A and KDM6B double deficient U937 and KDM6B deficient HL60 cells were also established.

### Lentiviral *shRNA* screening

TEX, a ‘stem cell-like’ human hematopoietic line, was generated by expressing TLS-ERG leukemia fusion oncogene in cord blood derived HSPCs (*34*). This leukemia model line maintains functional heterogeneity of LSCs, cytokine dependency, and a functional p53 pathway (*35*). TEX cells growing in the logarithmic phase were infected with a genome scale lentivirus library comprising ∼78,432 *shRNAs* targeting 16,056 unique RefSeq genes (*32, 33*). Cells were infected and puromycin resistant clones selected to ensure single *shRNA* clone integration per cell. The cells were subjected to low dose of γ-IR (1 Gy) and allowed to recover for 7-10 days. A part of the cell was harvested for genomic DNA isolation after single round of irradiation. The remaining cells were subjected to two more rounds of irradiation and recovery prior to genomic DNA isolation. Genomic DNA from d0 (non-irradiated), 1 Gy x 1 and 1 Gy x 3 irradiated cells were subjected to sequencing and relative abundance of *shRNA* clones in non-irradiated and treated samples were determined by deep sequencing analysis. Gene lists were generated based on the *shRNA* enrichment score from the screen and ratios were calculated [IR (treatment)/NT (control)] and assigned to each barcoded *shRNA* corresponding to a particular gene. A random index was assigned for each of the gene, and cut-offs of 1.0 and 0.1 were set respectively for the sensitizing and resistance tails. The scatter plots were generated using the ‘ggplot’ function in ‘R’ and the ‘candidate genes’ were labeled only if they qualify the cut-off criteria of 1.6 for ‘1 round of radiation-d10 recovery cycle’ or 1.2 for ‘3 rounds of radiation-d10 recovery cycles’ respectively.

### RNA-sequencing and analysis

Bulk RNA sequencing (RNA-seq) experiments (n=2-5) were performed at Core Technologies Research Initiative, National Institute of Biomedical Genomics, Kalyani, India and raw data was analyzed by Bionivid Technology, Bangalore, India (*11, 37, 68*). In addition, an independent set of bulk RNA-seq experiment (three biological replicates in each group from wild type or KDM6 deficient U937 cells) was also performed at The Princess Margaret Genomics Centre, UHN, Toronto. Total RNA was isolated from U937 cells expressing *sh-C* or *sh-KDM6A* or *sh-KDM6B* or both using TRIzol (Thermo Fisher Scientific). Genomic DNA contamination was removed by DNase I treatment of the isolated RNA using DNase I RNase free kit (Roche). After quantification of the RNA and quality check, equal amount of RNA from each sample or cell types were used to generate the library using the TruSeq RNA Sample Prep Kit v.2 (Illumina). Paired-end sequencing was performed using TruSeq 3000 4000 SBS Kit v.3 (Illumina) on the HiSeq 4000 platform (*11, 37*). Quality control using the NGS QC Toolkit yielded 85.94% high quality reads on average which was used for downstream analysis. Using TopHat platform 79.25% high quality 38 x 10^6^ reads on average mapped to the *Homo sapiens* (hg19) reference genome, thereby suggesting a good quality of RNA-seq. Using Cufflinks-based maximum likelihood method transcripts were given a score for their expression. 29,686 gene transcripts were identified to be expressed in either of the cells using Cuffdiff validation. These transcripts represented 12,078 genes. Characterization of the transcript type identified 95.5% ‘Full Length’ or ‘Known Transcripts’ and 4.5% ‘Potentially Novel Isoforms’ in accordance with Cufflinks Class Code distribution. This indicates a largely complete transcription machinery activity. Utilizing GO-Elite v.1.2.5 Software (http://www.genmapp.org/go_elite/) significant biology analysis for differentially expressed transcripts was performed. Significantly enriched GO pathways were determined keeping *P* value at < 0.05. In testing for differential expression, we considered log2 FC > +1 (up-regulation) and log2 FC < -1 (down-regulation).

The Gene Set Enrichment Analysis (GSEA) tool (*69*) was used to assess whether loss of KDM6A decreased enrichment for the DDR signature. Overall, 10,000 permuations of gene sets were used to compare the KDM6A deficient and control AML cases. *P*-values were corrected for family-wise error rate. The distribution of normalized enrichment scores was plotted as a boxplot. CD34^+^CD38^-^CD45RA^-^ HSCs were also isolated from pooled umbilical cord blood samples and subjected to 3 Gy irradiation at Tel Aviv University (*32, 33*). The cells were then allowed to recover for 4 hr and 24 hr. Control (non-irradiated) and irradiated cells were subjected to bulk RNA-seq to determine gene sets that were transcriptionally up or downregulated.

### ATAC-seq processing and analysis

Assay for Transposase-Accessible Chromatin with high-throughput sequencing (bulk ATAC-seq) experiments were performed in triplicate in each group of control or KDM6A and KDM6B deficient U937 cells at The Princess Margaret Genomics Centre, UHN, Toronto. Library preparation for ATAC–seq was performed on 5,000 cells with Nextera DNA Sample Preparation kit (Illumina), according to a previously reported protocol (*70, 71*). Four ATAC–seq libraries were sequenced for each lane in a HiSeq 2500 System (Illumina) to generate paired-end 50-bp reads. Reads were mapped against the hg19 human reference genome using BWA with default parameters. All duplicate reads, and reads mapped to mitochondria, chrY, an ENCODE blacklisted region or an unspecified contig, were removed (ENCODE Project Consortium 2012). MACS 2.0.10 was used to call peaks with a uniform extension size of 147 bp. A set of 500-bp windows over hg19 overlapping by 250 bp was generated. For each sample, the windows overlapping a called peak were identified. Windows were considered to be present in each condition if they overlapped a peak in two replicates. Windows identified as present in one condition were considered specific to that condition. Motif enrichment was performed using HOMER against a catalog of all peaks called in any hematopoietic population was produced by merging all called peaks that overlapped by at least one base pair using bedtools.

### Chromatin Immunoprecipitation-sequencing (ChIP-seq) and analysis

ChIP-seq experiments were carried out at Core Technologies Research Initiative (CoTeRI), National Institute of Biomedical Genomics (NIBMG), Kalyani, India (*11, 37*). For each ChIP set, 10 x 10^6^ primary AML BM nuclear cells, or 10 x 10^6^ wild type or KDM6 deficient U937 cells were crosslinked with formaldehyde (Millipore-Sigma) in culture media. After cross-linking, chromatin was extracted and sonicated to fragment lengths between 150 bp and 900 bp in chromatin extraction buffer containing 10 mM Tris pH 8.0, 1 mM EDTA pH8.0, and 0.5 mM EGTA pH8.0. Chromatin was incubated with antibodies to KDM6A (A302-374A, Bethyl Laboratories), H3K27ac (ab4729, Abcam), H3K27me3 (07-449, Millipore), p300 (A300-358A, Bethyl Laboratories), CBP (D6C5, Cell Signaling Technology), SMARCC1 [sc-9746 (R-18), Santa Cruz Biotechnology), CHD4 (clone 3F2/4, ab70469, Abcam), HDAC1 (A300-713A, Bethyl Laboratories), and rabbit IgG (clone P120-101, Bethyl Laboratories) or mouse IgG (clone G3A1, 5415S, Cell Signaling Technology) overnight at 4°C with rotation. All antibodies were used at 1:1000 dilution. Protein A/G agarose beads (Cell Signaling Technology) were then added and incubated for 2 hr at 4°C. The beads were washed with chromatin extraction buffer and by increasing salt concentration. The chromatin was eluted from the beads in chromatin elution buffer at 65°C with gentle vortexing. The eluted chromatin was treated with RNase for 30 min at 37°C. Reverse cross-linking was performed by treating the eluted chromatin with Proteinase K (Millipore-Sigma) at 65°C for 2 hr. The DNA was finally precipitated by phenol-chloroform extraction; precipitated DNA was dissolved in Tris-EDTA buffer and subjected to ChIP-seq analyses.

Size distribution of the ChIP-enriched DNA was checked using high-sensitivity chips in the 2100 Bioanalyzer (Agilent Technologies) for each sample, and quantitation was performed in the Qubit Fluorometer (Thermo Fisher Scientific) by the picogreen method. ChIP-seq library preparation was performed using the TruSeqChIP Sample Prep Kit (Illumina) according to the manufacturer’s instructions. 10 ng of input ChIP-enriched DNA was used for ChIP-seq library preparation. Final libraries were checked using high-sensitivity chips in the 2100 Bioanalyzer. The average fragment size of the final librarieswas found to be 280 ± 8 bp. Paired end sequencing (2 x 100 bp) of these libraries were performed in the HiSeq 2500 (Illumina). Quality control analysis of the raw data using the NGS QC ToolKit was done, and high-quality (HQ) reads with filter criteria of bases having ≥20 Phred score and reads with ≥70% were filtered. Paired end reads (.fastq format) were aligned with Bowtie software using –best and –m 2 [i.e., mismatches against reference genome Ensembl build GrCh37/hg19 (considering 2% input as the baseline)], and saved in SAM format, which was then converted to a sorted BAM file using SAMTools. PCR duplicates were removed using SAMTools rmdup. Peak calling was performed using MACS14 model building with a *P* value cutoff of 0.05. Annotation of the identified peaks was performed with PeakAnalyzer. Functional enrichment analysis [gene ontology (GO) and pathway] was done using the Database for Annotation, Visualization and Integrated Discovery (DAVID) v.6.8 (https://david.ncifcrf.gov/). The gene list was uploaded and converted to respective gene identifiers from the U.S. National Center for Biotechnology Information. The converted gene list was submitted to DAVID, and functional annotation clustering was carried out, which comprises GO and pathway analysis. The R bioconductor package ChiPseeker (https://guangchuangyu.github.io/software/ChIPseeker/) was used to generate heatmaps, average profile distibutions, and pie charts. Bigwig/bed files were imported into the Integrative Genomics Viewer (http://software.broadinstitute.org/software/igv/home), and snapshots of particular genomic loci were captured.

### Gene enrichment and functional annotation

GO analysis was carried out using The Database for Annotation, Visualization and Integrated Discovery (DAVID) v6.8. The *P* value used in the analysis is a modified one, termed as EASE score threshold (maximum probability). The threshold of EASE Score is a modified Fisher Exact *P* value used for gene-enrichment analysis. It ranges from 0 to 1. Fisher Exact *P* value=0 represents perfect enrichment. Usually *P* value is equal or smaller than 0.05 to be considered strongly enriched in the annotation categories.

### Quantitative reverse transcription and PCR (qRT-PCR)

Total RNA was isolated using TRIzol (Life Technologies) according to manufacturer’s recommendation. Genomic DNA contamination was removed using DNase I recombinant, RNase free kit (Roche). RNA amount was quantified and cDNA was prepared using TaqMan Reverse Transcription Reagents (Applied Biosystems). Gene expression levels were determined by quantitative PCR performed using cDNA with iTaq Universal SYBR Green Supermix (Biorad) on the 7500 Fast Real-Time PCR System (Applied Biosystems). GAPDH was used as a housekeeping gene. Relative expression levels were calculated using the 2^−ΔΔCt^ method (*4, 11*). qRT-PCR primer details are available in Table S2.

### ChIP followed by quantitative PCR (qChIP) analysis

For each ChIP set, 1% Formaldehyde (final concentration) was added to culture media for 20 min at room temperature to crosslink 5 x 10^6^ cells (*11, 37*). To quench the reaction glycine was added to the crosslinked cells at room temperature for 5 min. After crosslinking, chromatin was extracted and sonicated to fragment lengths between 150-900 bp in chromatin extraction buffer containing 10 mM Tris pH=8.0, 1 mM EDTA pH=8.0 and 0.5 mM EGTA pH=8.0. After sonication, chromatin was clarified by centrifugation at 10,000 rpm in a microcentrifuge for 10 min at 4°C and analyzed for fragmentation size prior to immunoprecipitation. The chromatin was then incubated with 5 µg of antibody against KDM6A (A302-374A, Bethyl Laboratories and 33510S, Cell Signaling Technology), KDM6B (ab169197, Abcam), p300 (A300-358A, Bethyl Laboratories), CBP (clone D6C5, 7389S, Cell Signaling Technology), SMARCC1 (clone R-18, sc-9746, Santa Cruz Biotechnology), H3K27ac (ab4729, Abcam), H3K27me3 (07-449, Millipore), CHD4 (clone 3F2/4, ab70469, Abcam), HDAC1 (A300-713A, Bethyl Laboratories), and rabbit IgG (clone P120-101, Bethyl Laboratories) or mouse IgG (clone G3A1, 5415S, Cell Signaling Technology). ChIP-ed DNA or chromatin was purified using phenol-chloroform extraction and chromatin occupancy was determined using primers designed for TSS and upstream promoter regions (−0.5 Kb) of target gene loci. For phenol-chloroform extraction, sample volume was increased to 300 μL by adding TE and 7 μL of 5M NaCl. To the mixture an equal volume (300 μL) of phenol-chloroform and isoamyl alcohol was added. The tubes were vortexed and centrifuged at 14000 rpm for 5 min at room temperature to collect the aqueous phase. This was repeated twice. Further 300 μL of chloroform was added to the aqueous phase and the tubes vortexed vigorously. The samples were centrifuged at 14,000 rpm for 5 min at room temperature. DNA was precipitated overnight at -20°C using ethanol. Purified DNA was pelleted down by centrifugation at 14,000 rpm for 15 min at 4°C and washed with 75% ethanol. The pellet was air dried completely and redissolved in 20-30 μL of TE or DNase-free water. qChIP primer details are available in Table S3.

### Drug treatments and cell viability assays

AML cells were cultured in IMDM or RPMI or α-MEM media (depending on cell type) supplemented with 10% or 20% FBS, 100 U/mL Penicillin, 100 μg/mL Streptomycin and 2 mM L-Glutamine (all from Gibco) at 37°C with 5% CO_2_. GSK-J4 was purchased from Sigma Aldrich (SML0701), and olaparib (10621) and venetoclax (16233) were procured from Cayman Chemicals. For IC_50_ analysis, 50,000 AML cells were seeded in 200 µL complete media in 48 well non-TC plates (BD Falcon), and treated with varying doses (from 1 nM to 1 mM) of GSK-J4 or olaparib or venetoclax for 72 hr. Drug combination assays were performed using 1:1 ratios simultaneously, and trypan blue negative viable cell counts were determined after 48 hr to 72 hr of treatment with individual drugs or in combinations. Combination index (Ci) at ED_50_ was calculated using CompuSyn v 1.0. Ci < 1 was considered as drug synergism. In addition, in some of the experiments a constant dose of GSK-J4 was also used, set at half of the IC_50_ concentrations of respective cell types, to analyze GSK-J4 combination with venetoclax or AraC or daunorubicin. For analyzing BCL2 expression, AML cells were treated with GSK-J4 for 24 hr to 72 hr at the IC_50_ concentrations of respective cell types.

In separate experiments, control or KDM6A deficient U937 cells were transduced with lentiviral particles expressing *FLT3-ITD*. After 48 hr of transduction, the cells were treated with olaparib with concentrations between 1 μM to 100 μM. Niraparib tosylate (MK-4827) and navitoclax (ABT-263) were purchased from Cayman Chemicals. For calculating IC_50_ of niraparib, OCI-AML-2 and OCI-AML-5 cells were subjected to a range of concentration between 1 μM to 100 μM. Similarly, navitoclax IC_50_ was calculated by exposing the cells to drug concentrations ranging from 1 nM to 50 μM. For survival assay, AML cells were cultured in presence of GSK-J4, olaparib, venetoclax or DMSO (vehicle) alone or in combination for 48 hr. *KDM6A* mutant and *KDM6A* wild type primary AML peripheral blood mononuclear cells were used to study *in vitro* efficacy of olaparib and venetoclax combination therapy.

### γ-irradiation and PARP inhibition assays

For DSB induction, AML cells were irradiated with low dose 3 Gy or lethal 10 Gy γ-IR using ^60^Co Blood Irradiator (1.485 Gy/min) at CSIR-Indian Institute of Chemical Biology gamma irradiation core facility. Cells were then allowed to recover upto 8 hr followed by staining with antibodies for flow cytometry analysis. For proliferation assays, wild type or KDM6 deficient AML cells were allowed to grow in triplicate in regular media in presence of 10 µM of olaparib. Trypan blue-negative cell counts were determined at different time points. For apoptosis assay, AML cells were cultured for 72 hr in presence of 50 µM olaparib or DMSO (vehicle).

### HR and NHEJ reporter assay

To determine HR-NHEJ activity leukemia cells were subjected to GSK-J4 (2 μM) or DMSO (vehicle) treatment for 16 hr. Following treatment 1-2 x 10^6^ cells were nucleofected or transfected with 5 µg of I-SceI-linearized HR or NHEJ reporter plasmid and 2.5 µg of mCherry plasmid (transfection efficiency control) (*61*). Nucleofector program FF-120, SF Cell Line 4D-Nucleofector™ x Kit (Lonza) were used. Repair of linearized plasmid by HR or D-NHEJ event allowed GFP expression. After 72 hr, the percentage of GFP^+^ cells in mCherry^+^ cells were analyzed by flow cytometry to assess HR and NHEJ activity.

### Comet assay

DNA damage was compared using neutral comet assay (*72*). Briefly, after γ-irradiation treatment (10 Gy), cells were incubated for 2 hr at 37^0^C and 5% CO_2_. Next the cells were mixed with low melting agarose (Sigma) and spread on a pre-warmed glass slide. The slides were immersed in lysis solution (2.5 M NaCl, 100 mM disodium EDTA, 10 mM Tris base, 200 mM NaOH, 1% lauryl sarcosinate and 1% Triton-X 100, pH 10, all from Sigma) at 4°C overnight, rinsed with deionized water, and immersed in a 4°C neutral electrophoresis solution (100 mM Tris base and 300 mM sodium acetate, adjusting the pH to 9.0 with glacial acetic acid) for 1 hr in dark. Slides were subjected to electrophoresis at a constant voltage of 1V/cm for 30 min at 4°C. Thereafter, the slides were immersed in DNA precipitation solution (7.5 M ammonium acetate, with 95% ethanol) for 30 min at room temperature and washed with 70% ethanol for 30 min. DNA was stained with SYBR Green (Invitrogen), and images were captured in fluorescence microscope (EVOS Cell Imaging System, Invitrogen). Comet Score 2.0 software was used to measure comet tail length for about 100 cells. Statistical analysis of comet tail lengths was performed using the student’s *t*-test.

### γH2A.X staining and immunofluorescence analysis

AML cells were subjected towards 10 Gy of γ-IR and allowed to repair for 15 min and 2 hr. Cells were fixed with 1.5% paraformaldehyde (Sigma) for 15 min at room temperature and permeabilized with 0.1 % Triton X-100 (Sigma) for 10 min. After washing with PBS, cells were blocked using 1% BSA and stained with anti-γH2A.X antibody (Millipore, 05-636) at 1:100 dilution for overnight at 4°C, which was followed by incubation with secondary antibody Dy-light 488 (Bethyl, A21202) for 1 hr in room temperature. The cells were washed three times with PBS and cytocentrifuged (CellSpin, Hanil Scientific) at 700 g for 10 min at room temperature. Slides were mounted with ProLong Gold antifade reagent with DAPI (Invitrogen). Images were captured using confocal microscope (ZEISS, LSM 980) in 200X resolution and number of γH2A.X foci per nucleus was quantified using ZEISS ZEN 3.5 (ZEN lite) software.

### Flow cytometry and BCL2 expression analysis

AML cells were assessed by flow cytometry using anti-H3K27me3 (Millipore), anti-KDM6A (Bethyl) and anti-BCL2 (15071T, Cell Signaling Technology, 1:200 dilution). All antibodies were used at a dilution of 1:100. For immunophenotyping anti-human CD34-PE and anti-human CD45-PECy7 (BD Biosciences) antibodies were used at 1:20 dilution. Briefly, using 1.5% paraformaldehyde (Sigma) 1-2 x10^6^ cells were fixed and then permeabilized with methanol. After fixation the cells were stained for histone modifications or KDM6A, washed with PBS supplemented with 2% human serum, and incubated with anti-rabbit IgG-DyLight488 (Bethyl Laboratories) for 10 min in dark. For immunophenotyping, the cells were incubated with fluorescent-dye labelled antibody during secondary antibody incubation. For γH2A.X and p-ATM analysis, after washing the cells with ice cold PBS, cells were fixed using 70% ethanol keeping cell density at 1 x10^6^ cells /mL. Anti-γH2A.X specific antibody (Millipore) or anti-pATM specific antibody (Cell Signaling Technology) was added at 1:100 dilution to the cells for overnight incubation at 4°C, followed by secondary antibody incubation for 1 hr at room temperature. To remove any non-specific antibody binding cells were washed with PBS and re-suspended in 500 µL of PBS supplemented with 2% human serum. Cells were analyzed using LSRFortessa (Becton Dickinson) and FACSDiva software (Becton Dickinson). Flow Jo v10 was used to make univariate staggered histogram plots for mean fluorescence intensity analysis.

### ROS staining

Cells were treated with GSK-J4 (2 μM) or olaparib (10 μM) individually or in combination and DMSO (vehicle) for 72 hours. Following treatment CellROX Reagent (Thermo Scientific) was added at a final concentration of 5 μM to the cells and incubated for 30 min at 37°C. Cells were washed three times with PBS. 7-AAD was then added to the cells to determine the ROS status in live cells. Cells were analyzed by flow cytometry in LSRFortessa (Becton Dickinson) using FACSDiva software (Becton Dickinson).

### Mito-Red and Mito-Green staining

AML cells were treated with GSK-J4 (2 μM) or olaparib (10 μM) individually or in combination, and DMSO (vehicle) for 72 hr. Stock solution of MitoTracker Red CMXRos (Thermo Scientific) and MitoTracker Green FM (Thermo Scientific) were prepared in DMSO to a concentration of 1 mM. Prior to use the stock solution was diluted to a final working concentration of 50 nM for MitoTracker Green FM and 100 nM for MitoTracker Red CMXRos. While MitoTracker Green FM binds to mitochondria irrespective of mitochondrial membrane potential, MitoTracker Red CMXRos stains mitochondria in live cells and its accumulation is dependent upon membrane potential. Briefly, cells were harvested by centrifugation and resuspended gently in pre-warmed (37°C) staining solution containing the MitoTracker probe. Cells were incubated for 30 min at 37°C. Post staining cells were washed twice in PBS and redissolved in FACS buffer prior to analysis in LSRFortessa (Becton Dickinson).

### PARP inhibition assay

About 1 x 10^6^ U937 and OCI-AML-5 cells were treated with DMSO or olaparib (1 µM) for 48 hr. Following treatment the cells were washed twice with PBS, and fixed in 4% formaldehyde for 20 min at room temperature. Cells were then permeabilized with 0.1% Triton X-100 for 15 min. After fixation the cells were stained with anti-Poly-(ADP-ribose) polymer antibody (Abcam, ab14459) for 1 hr at 37°C. Cells were washed and stained with secondary antibody for 1 hr at room temperature. To remove any non-specific antibody binding cells were washed with PBS and re-suspended in 500 µL of PBS supplemented with 2% human serum. The cells were analyzed using LSRFortessa (Becton Dickinson) and FACSDiva software (Becton Dickinson).

### Apoptosis assay

For apoptosis assay cells were washed twice with ice cold PBS and resuspended in 1x Annexin V binding buffer keeping concentration at 1 x 10^6^ cells/mL. Annexin V APC (1:20) (BD PharMingen) and 7-AAD (1 µg/mL) were added to the tubes. The cells are vortexed gently and incubated at room temperature in the dark for 15 min. After incubation, 200-400 μL of 1x Binding Buffer was added to each tube and analyzed in a LSRFortessa (Becton Dickinson) using FACSDiva software (Becton Dickinson) within 30 min.

### Immunoblot analysis

Total cell lysates were prepared by incubating cells in 1x RIPA (Cell Signaling) containing protease and phosphatase inhibitor cocktails for 15 min followed by brief sonication. Supernatants were collected after centrifugation at 16,000 g for 15 min at 4^0^C. Protein concentration was determined using Pierce BCA Protein assay kit (Thermo), and lysates were resuspended in 1x SDS gel loading buffer. The proteins were separated in SDS-PAGE and transferred to PVDF membrane (Millipore) and subsequently probed with respective antibodies including anti-RAD51 (8875S, Cell Signaling Technology), anti-RAD50 (3427T, Cell Signaling Technology). All antibodies were used at a dilution of 1:1000 unless specified otherwise.

### Histone extraction and immunoblot analysis

Total histone was isolated from wild type or KDM6A and/or KDM6B deficient U937 cells using EpiQuik total histone isolation kit (Epigentek) according to manufacturer’s protocol. Briefly 3 x10^6^ cells were harvested and incubated in 1x pre-lysis buffer for 10 min in ice. Following incubation, the tubes were centrifuged and supernatant discarded. The pellet was resuspended in lysis buffer and incubated in ice for 30 min. The supernatant containing acid soluble protein was collected in a fresh tube and neutralized using Balance-DTT buffer. Protein concentration was determined using BCA protein assay kit (Pierce). Lysates were resuspended in 1x SDS gel loading buffer, and the proteins were separated in SDS-PAGE and transferred to PVDF membrane (Millipore) and subsequently probed with respective antibodies.

### Analysis of OHSU AML cohort

Heatmap cluster of KDM6A, KDM6B and PARP1 expression from OHSU AML cohort was derived using cBioPortal for Cancer Genomics interface. For correlation analysis, mRNA expression (RNA Seq v.2 RSEM) was obtained and plotted using GraphPad Prism 5 v5.0.

### Statistical analyses

Statistical analyses were performed using GraphPad Prism v 8.0.2. Student’s t-test was used to determine statistical significance, using Welch’s Correction wherever the variances were significantly different. To perform multiple comparisons of data that is normally distributed and have equal variance, one way ANOVA was used followed by Tukey Kramer post hoc test. Quantitative results are expressed as means ± SD or SEM. For IC_50_ calculation, cell counts were normalized with respect to minimal drug concentration as control and plotted against logarithm of respective drug concentrations. For all the cell viability plots, nonlinear regression curves were obtained between logarithms of respective drug concentration and normalized viable cell count. Drug combination index (Ci) at ED_50_ was calculated using CompuSyn v 1.0. Ci < 1 was considered as drug synergism. Flow Jo v10 was used to make univariate staggered histogram plots for mean fluorescence intensity analysis. Densitometry analyses were performed using NIH ImageJ. For all statistical analyses, the level of significance was set at 0.05.

## Acknowledgments

We thank Drs. Kristian Helin (Addgene# 24168), Kai Ge (Addgene# 40619), Didier Trono (Addgene# 12260, 12259), Tomasz Skorski and Steven Chan for sharing plasmid constructs and cells. We also thank Alexander Avgoustis for helping with AML specimens through Leukemia Tissue Bank at Princess Margaret Cancer Centre/University Health Network. We acknowledge Nicholas Khuu, Julissa Tsao (Princess Margaret Genomics Centre, Toronto), and Drs. Arindam Maitra, Disha Banerjee, Subrata Patra (National Genomics Core/Co-TERI, National Institute of Biomedical Genomics, Kalyani, India) for next-generation sequencing services, Princess Margaret (UHN) Animal Resources Centre, Flow Cytometry Core and CSIR-IICB Flow Cytometry, Central Instrumentation, Radiation Facility and Dr. Arunima Maiti, Tata Translational Cancer Research Center (TTCRC) for Flow Cytometry sorting facility and experimental help. We thank Dr. Nabanita Dasgupta, NRS Medical College & Hospital for providing umbilical cord blood samples. We are grateful to Dr. Craig Jordan (U Colorado) for sharing transcriptome datasets of venetoclax-resistant Mono-AML, and we appreciate his comments during the preparation of this manuscript. We also thank Drs. Stephanie Xie, Helena Boutzen, Jean Wang and other members of the Sengupta and Dick laboratories for comments and helpful discussions, and Sally Desilva, Monica Doedens for administrative assistance.

## Authors contributions

Conception, study design and interpretation: A.S. Experimental design, data acquisition, analysis and interpretation: L.D.B., S.G., S.K.B. Immunoblot analysis: W.S., S.S.C. Biochemical studies: S.B., M.B., S.S., A.B., S.C. Xenotransplantation and pharmacological studies: L.J., N.M., O.I.G., A.S. Drug sensitivity and cell survival assays: S.G., S.K.B. Drug combination index analysis: W.S., S.B. ATAC-seq, RNA-seq and computational analysis: A. Mu. RNAi screening: S.S.N.A.M., M.M. Bioinformatics analysis: L.D.B., S.G., A.G.X.Z., M.B. AML tissue banking and characterization: A.A., J.A.K., A. Mi., E.R.L., D.B., M.D.M. Manuscript writing: L.D.B., A.S. Research direction, resources, fund acquisition, manuscript editing and overall supervision: J.E.D., A.S. All authors have contributed and agreed with the final version of the manuscript.

## SUPPLEMENTARY MATERIALS

**Fig. S1.**
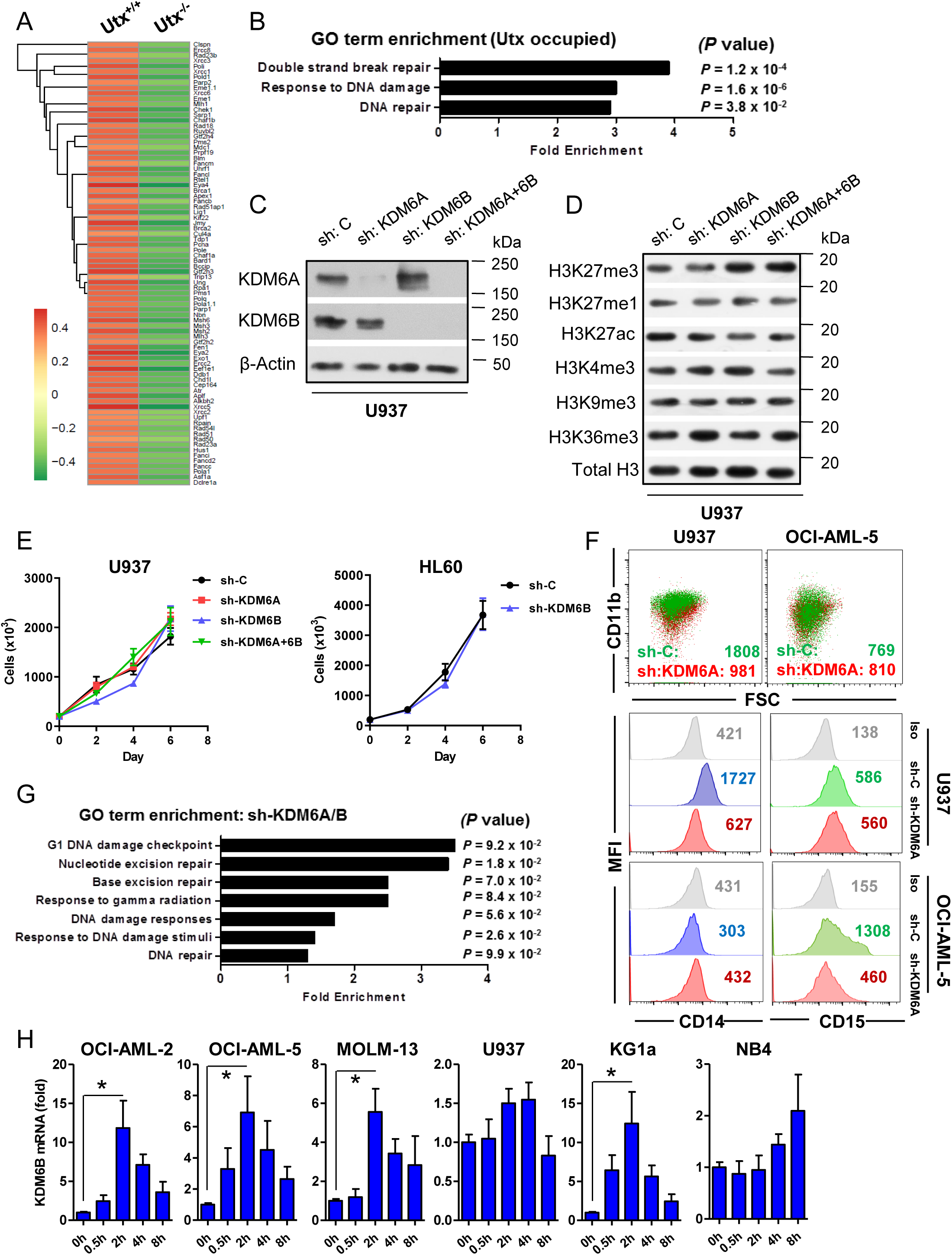
KDM6 expression in AML associates with DDR gene regulation. (A) RNA-seq heatmap analysis showing differential expression of DNA damage repair (DDR) pathway genes in *Utx^+/+^* Lin^-^ bone marrow (n=2) versus *Utx^-/-^*splenocytes (n=3). (B) GO term analysis of the 1825 downregulated genes occupied by Utx. (C) Immunoblot analysis in KDM6A and/or KDM6B deficient U937 cells. (D) Immunoblot analysis showing histone modifications in KDM6 deficient U937 cells. (E) Basal proliferation of control or KDM6 deficient AML cells. (F) Flow cytometry dot plots *(upper)* and histogram analysis *(lower)* in control or KDM6A deficient AML cells at steady state. (G) GO term analysis of the 2272 downregulated genes in KDM6 deficient U937 cells compared to control. (H) qRT-PCR (normalized to 0h) of KDM6B in AML cells treated with 10 Gy of γ-IR (n=3). qRT-PCR values were normalized to GAPDH. Statistics were calculated with Student’s t-test; error bars represent means ± SD. \**P* < 0.05 was considered to be statistically significant.

**Fig. S2.**
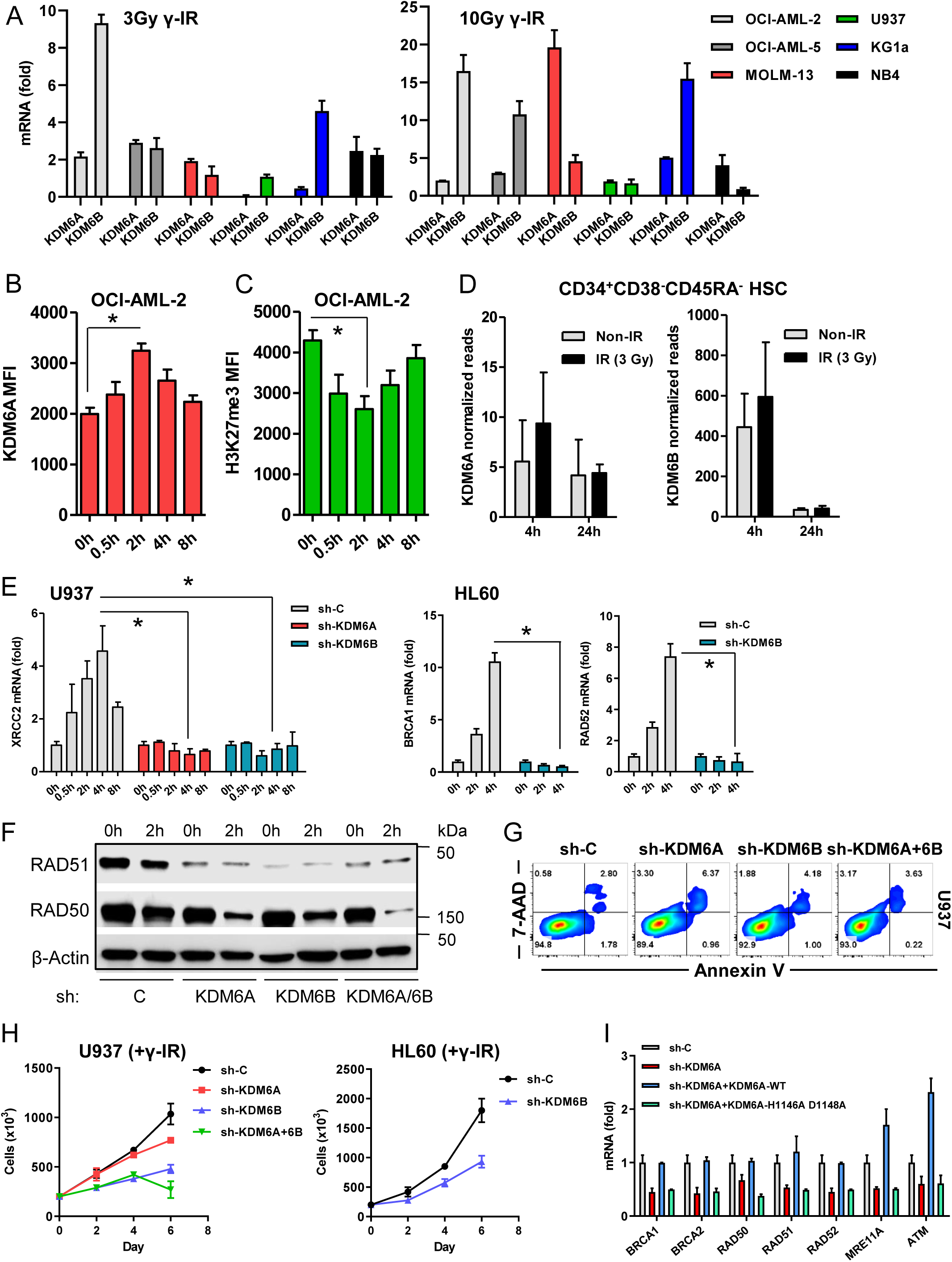
KDM6 demethylases correlate with DDR in AML. (A) qRT-PCR analysis (normalized to 0h) in AML cells treated with 3 Gy *(left)* or 10 Gy *(right)* of γ-IR. (B) Flow cytometry analysis showing MFI of KDM6A in OCI-AML-2 cells irradiated with 10 Gy of γ-IR (n=3). (C) Flow cytometry analysis showing MFI of H3K27me3 in OCI-AML-2 cells irradiated with 10 Gy of γ-IR (n=3). (D) qRT-PCR analysis of KDM6A *(left)* and KDM6B *(right)* performed in cord blood derived CD34^+^CD38^-^CD45RA^-^ normal HSCs in response to 3 Gy of γ-IR (n=4). (E) qRT-PCR (normalized to 0h) analysis in control or KDM6 deficient AML cells treated with 10 Gy of γ-IR. (F) Immunoblot analysis performed in control or KDM6 deficient U937 cells treated with 10 Gy of γ-IR. (G) Flow cytometry analysis representing Annexin V^+^ control or KDM6 deficient U937 cells treated with 10 Gy of γ-IR. (H) Proliferation of control or KDM6 deficient AML cells in response to 10 Gy of γ-IR. (I) qRT-PCR analysis HR and MRN complex genes in control or KDM6A deficient 293T cells overexpressing either wild type *KDM6A* or demethylase-dead *KDM6A* point mutant. qRT-PCR values were normalized to GAPDH. Data are representative of three independent experiments. Statistics were calculated with Student’s t-test; error bars represent means ± SD. \**P* < 0.05 was considered to be statistically significant.

**Fig. S3.**
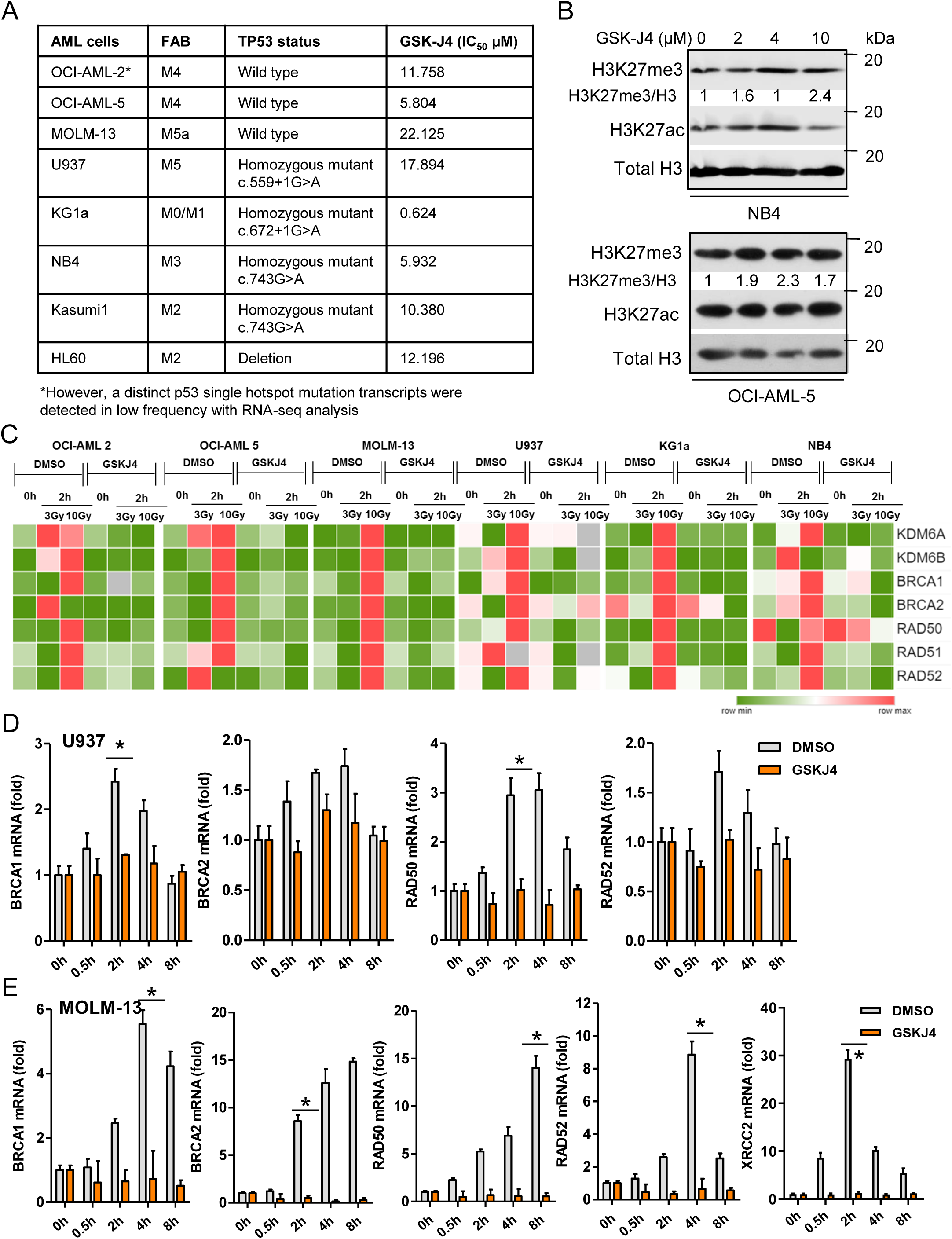
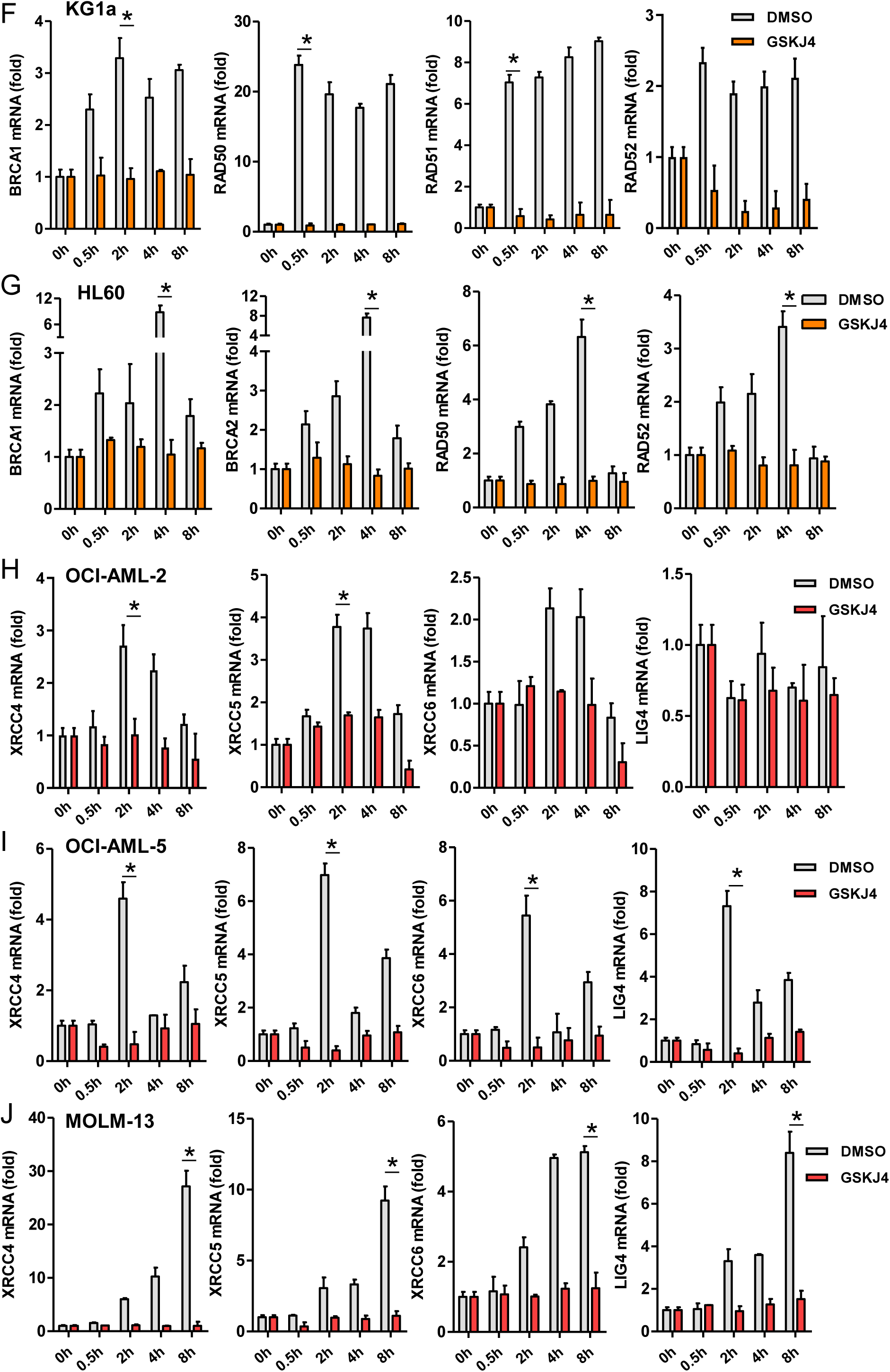
KDM6 inhibition impairs DDR gene expression in AML. (A) IC_50_ of GSK-J4 (treated for 72 hr) in various AML cells. (B) Representative immunoblot analysis showing histone modifications of acid extracted fractions of AML cells treated with varying concentrations of GSK-J4 for 48 hr. Normalized H3K27me3 levels are shown. (C) qRT-PCR (normalized to 0h) based heatmap analysis showing gene expression alterations in DMSO or GSK-J4 treated AML cells irradiated with 0 Gy, 3 Gy or 10 Gy of γ-IR. (D-J) qRT-PCR analysis (normalized to 0h) of HR genes in DMSO or GSK-J4 treated various AML cell lines in response to 10 Gy of γ-IR (n=2). qRT-PCR values were normalized to GAPDH. Data represent average of two to three independent experiments. Statistics were calculated with Student’s t-test; error bars represent means ± SD. \**P* < 0.05 was considered to be statistically significant.

**Fig. S4.**
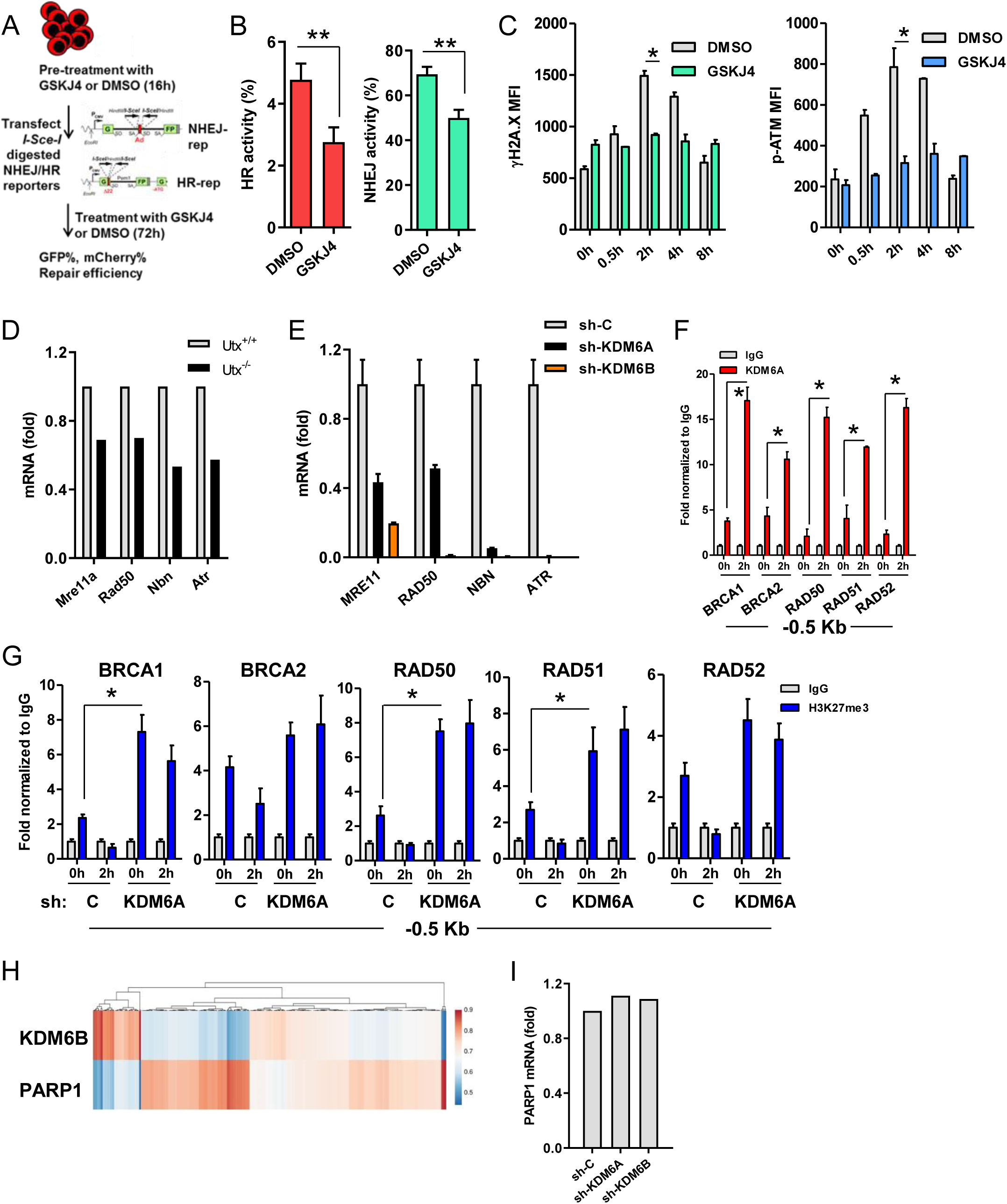
KDM6A regulates DDR transcriptional activation. (A) Schema representing assay for measuring repair efficiency using HR and NHEJ reporter constructs. (B) Analysis of HR *(left)* and NHEJ *(right)* activity in K562 cells using flow cytometry (n=3). Notorious co-transfection efficiencies hindered similar analysis in AML cells. (C) Flow cytometry analysis showing MFI of γH2A.X *(left)* and p-ATM *(right)* in DMSO and GSK-J4 treated U937 cells treated with 10 Gy of γ-IR (n=2). (D) RNA-seq mean expression of MRN complex subunits between *Utx^+/+^* (n=2) and *Utx^-/-^* (n=3) hematopoietic cells. (E) qRT-PCR of MRN in control or KDM6 deficient U937 cells. (F) qChIP analysis showing chromatin occupancy of KDM6A on HR gene promoter regions (−0.5 Kb) in U937 cells treated with 10 Gy of γ-IR. (G) qChIP showing chromatin occupancy of H3K27me3 on HR gene promoter regions (−0.5 Kb) in control or KDM6A deficient U937 cells in response to 10 Gy of γ-IR. (H) KDM6B and PARP1 mRNA expression z-scores (RNASeq v2 RSEM) heatmap cluster from OHSU AML dataset. (I) RNA-seq mean expression of PARP1 in control or KDM6A or KDM6B deficient U937. qChIP values were normalized to IgG. Data are representative of two independent experiments. Statistics were calculated with Student’s t-test; error bars represent means ± SD. \**P* < 0.05 or \*\**P* < 0.01 were considered to be statistically significant.

**Fig. S5.**
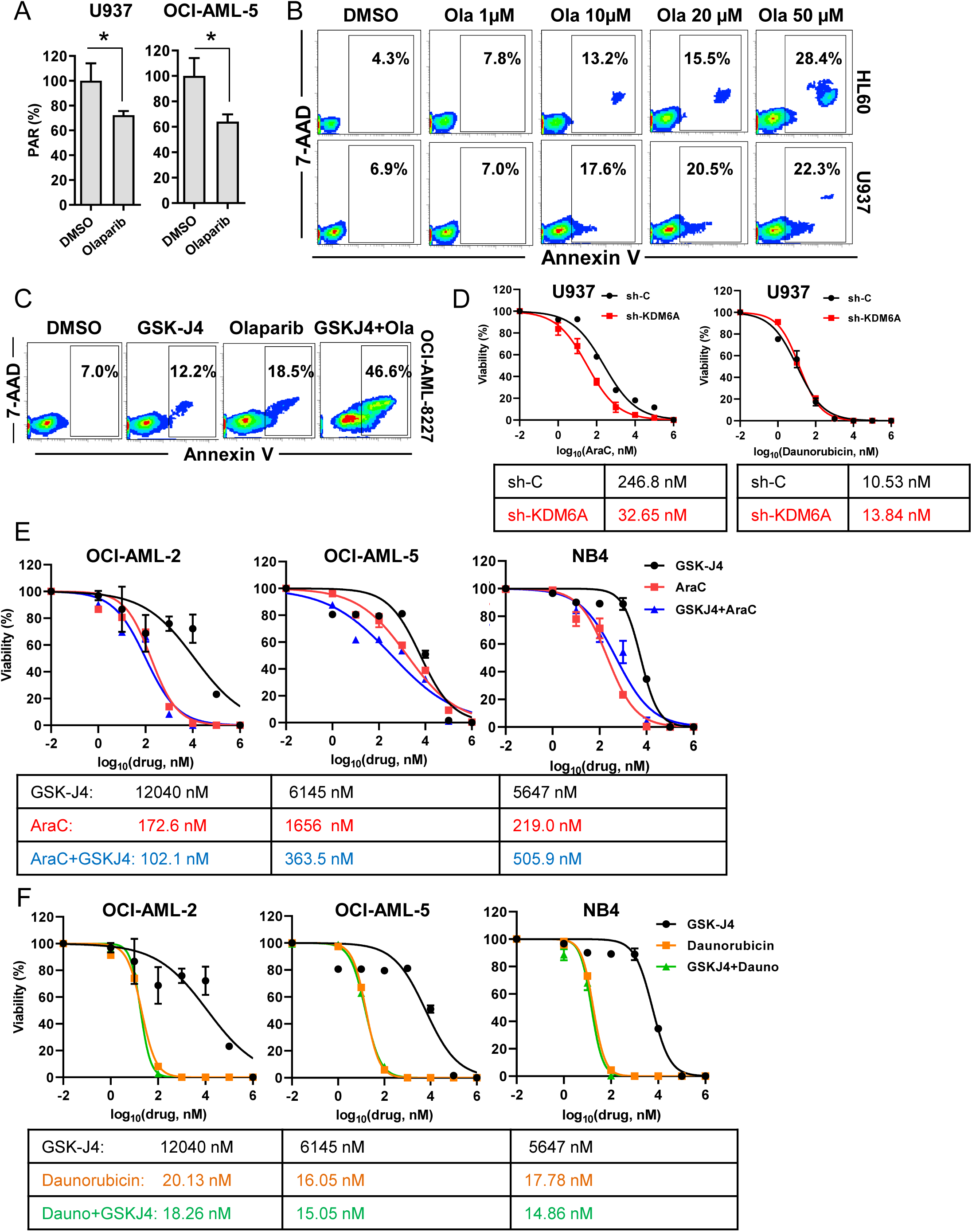
KDM6 deficient AML cells are sensitive to targeted therapy. (A) Flow cytometry quantitation of intracellular PAR levels in AML cells treated with DMSO or olaparib (1 µM). (B) Flow cytometry contour plots representing apoptotic cell populations in AML cells treated with increasing concentrations of olaparib. (C) Flow cytometry contour plots showing apoptosis in OCI-AML-8227 cells treated with GSK-J4 and/or olaparib. (D) Percent viability of control or KDM6A deficient AML cells treated with a varying doses (from 1 nM to 1 mM) of AraC *(left)* or daunorubicin *(right)* for 48 hr. Data represent average of two to three independent experiments with similar results. IC_50_ values are tabulated. (E) Percent viability of AML cells treated with varying doses (from 1 nM to 1 mM) of GSK-J4 alone *(black)* or AraC alone *(red)* or in combination *(blue)* for 72 hr. GSK-J4 dose in the combination dose is set at half of the IC_50_ values of respective cell types. Data represent average of two independent experiments. IC_50_ values are tabulated. (F) Percent viability of AML cells treated with varying doses (from 1 nM to 1 mM) of GSK-J4 alone *(black)* or daunorubicin alone *(orange)* or in combination *(green)* for 72 hr. GSK-J4 dose in the combination dose is set at half of the IC_50_ values of respective cell types. Data represent average of two independent experiments with similar results. Data represent average of two independent experiments. IC_50_ values are tabulated.

**Fig. S6.**
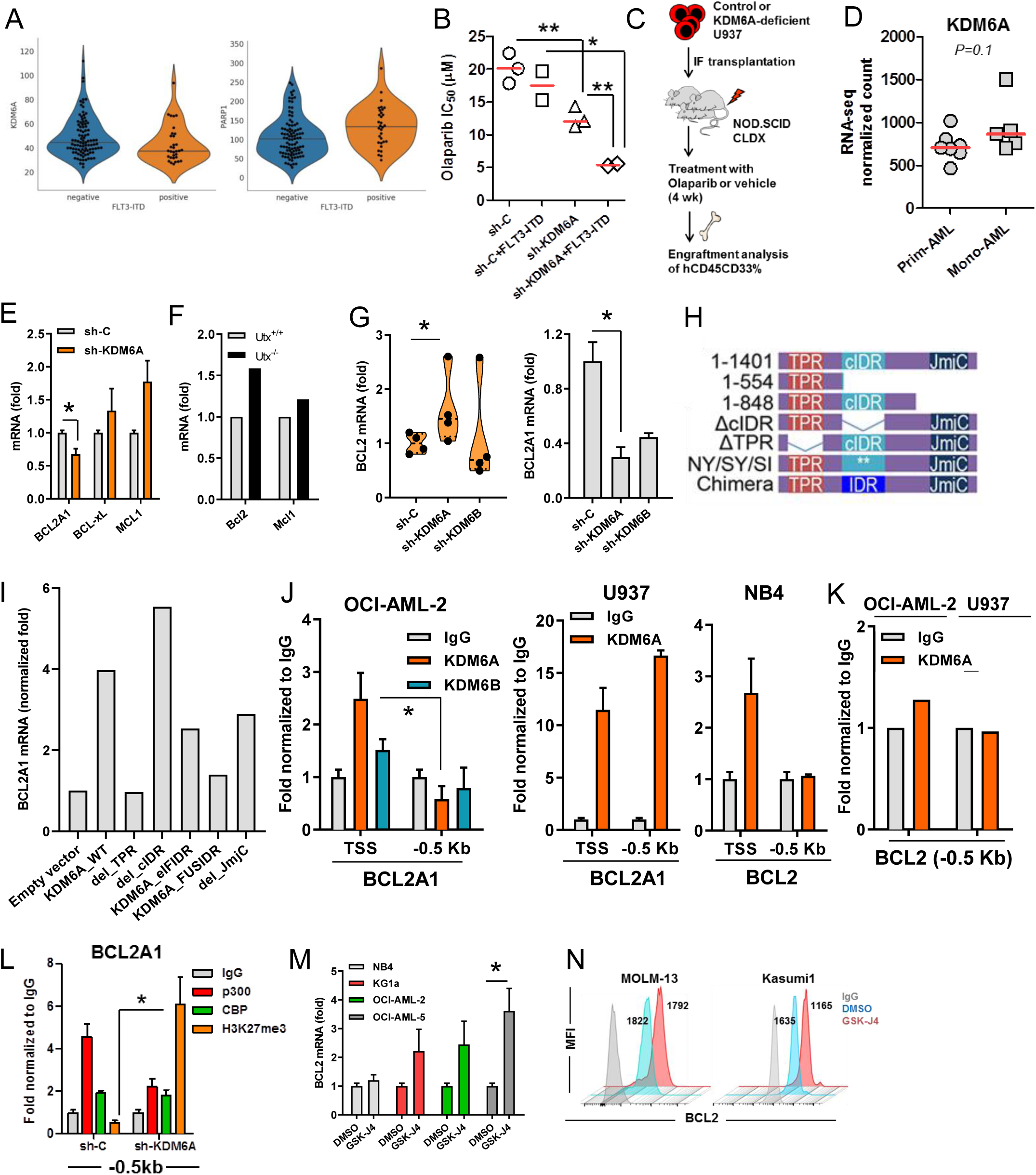
KDM6A associates with BCL2 family gene regulation and venetoclax sensitivity. (A) *FLT3-ITD* correlation with KDM6A *(left)* and PARP1 *(right)* expression from Beat AML (n=702). (B) IC_50_ of olaparib treated for 72 hr in control or KDMA deficient U937 expressing *FLT3-ITD*. (C) Schema representing bone marrow engraftment analysis performed in U937 cell lines-derived (control or KDM6A deficient) AML xenografts (CLDX) in response to PARP inhibition. (D) RNA-seq analysis showing expression of KDM6A between primitive (Prim-) AML (n=7) and monocytic (Mono-) AML (n=5) ROS^low^ LSCs. (E) qRT-PCR analysis of control or KDM6A deficient U937 cells. (F) RNA-seq mean expression in *Utx^+/+^* (n=2) and *Utx^-/-^* (n=3) hematopoietic cells. (G) RNA-seq analysis of BCL2 *(left)* and BCL2A1 *(right)* expression in control or KDM6A or KDM6B deficient U937 cells. (H) Architecture of KDM6A-domain mutants (adapted from B. Shi et al, Nature 2021). (I) RNA-seq mean expression analysis in KDM6A-null THP1 cells expressing full length KDM6A or various domain mutants. (J-K) qChIP analysis showing chromatin occupancy of KDM6 proteins on BCL2A1 and BCL2 in different AML cells. (L) qChIP analysis showing chromatin occupancy at BCL2A1 promoter region (−0.5 Kb) in control and KDM6A deficient U937. (M) qRT-PCR analysis of AML cells treated with GSK-J4 at half of the respective IC_50_ concentrations. Data represent average of four to five independent experiments; error bars represent means ± SEM. (N) Flow cytometry staggered histogram plots showing BCL2 expression in AML cells treated with DMSO *(blue)* or GSK-J4 *(red)* at respective IC_50_ concentrations for 48 hr. qChIP values were normalized to IgG. Data represent average of two to three independent experiments. Statistics were calculated with Student’s t-test; error bars represent means ± SD. \**P* < 0.05 or \*\**P* < 0.01 were considered to be statistically significant.

**Fig. S7.**
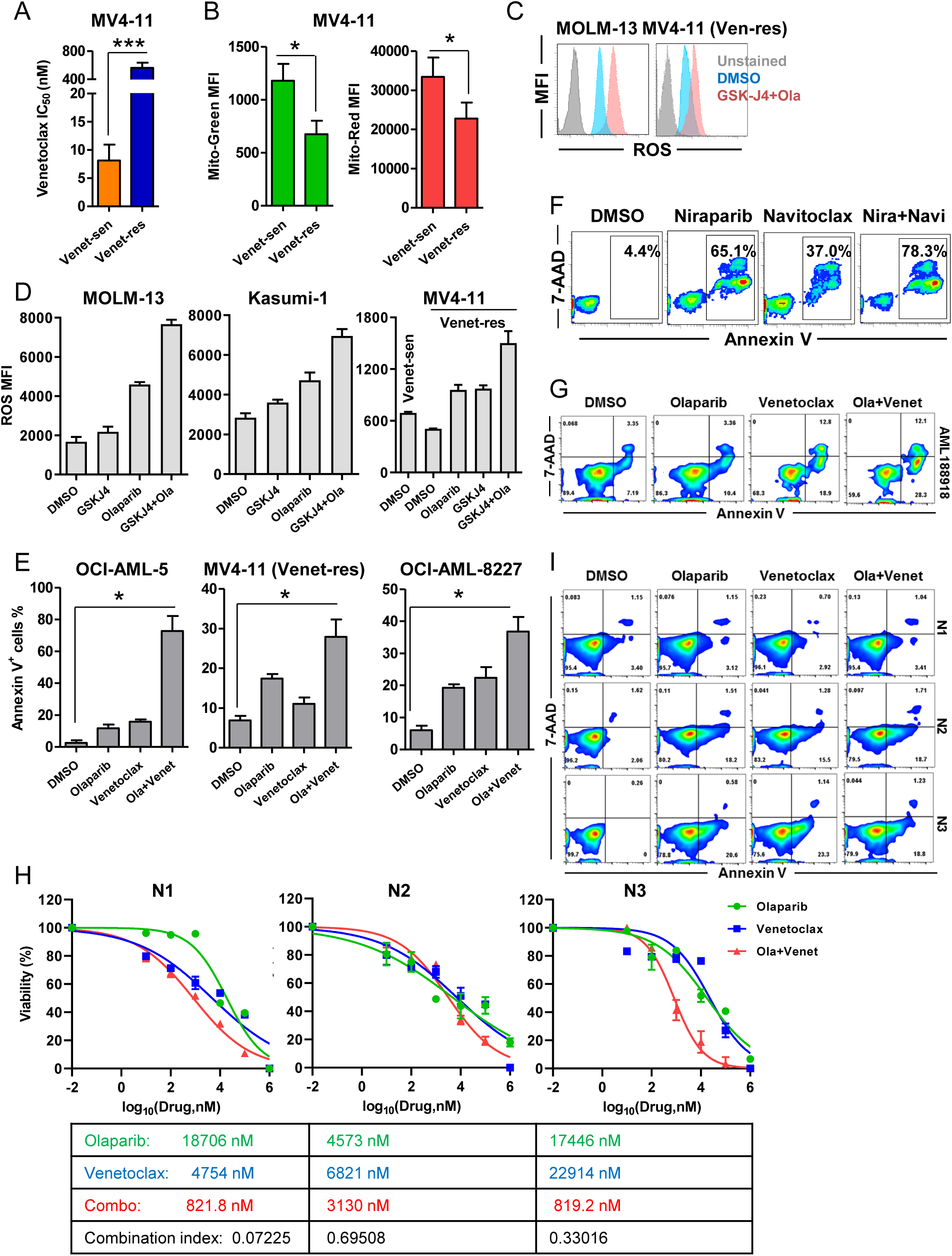
Combined inhibition of PARP and BCL2 in AML. (A) IC_50_ of venetoclax in venetoclax sensitive (Ven-sen) and venetoclax resistant (Ven-res) MV4-11 cells treated for 72 hr (n=3). (B) Flow cytometry analysis of mitochondrial mass (Mito-Green) *(left)* and mitochondrial membrane potential (Mito-Red) *(right)* in Ven-sen and Ven-res MV4-11 (n=3). (C) Flow cytometry histogram plots showing intracellular ROS in AML cells treated with DMSO *(blue)* or a combination of GSK-J4 and olaparib *(red)* for 48 hr. (D) Flow cytometry MFI analysis of intracellular ROS in AML cells treated with either DMSO, GSK-J4, olaparib, or a combination of GSK-J4 and olaparib for 48 hr. (E) Flow cytometry analysis showing apoptosis in AML cells treated with either DMSO, olaparib, venetoclax, or a combination of olaparib and venetoclax for 48 hr (n=2). (F) Flow cytometry analysis showing apoptosis in AML cells treated with either DMSO, niraparib, navitoclax, or their combination for 48 hr (n=2). (G) Flow cytometry analysis showing apoptosis of KDM6A-mutant AML188918 treated with either DMSO, olaparib, venetoclax, or a combination of olaparib and venetoclax at respective IC_50_ concentrations for 48 hr. (H) Percent viability of cord blood derived normal (N) CD34^+^ hematopoietic stem/progenitor cells (HSPCs) (n=3) treated with varying doses (from 10 nM to 1 mM) of olaparib alone *(green)* or venetoclax alone *(blue)* or in combination *(red)* for 48 hr. IC_50_ values are tabulated and Ci at ED_50_ was calculated using CompuSyn v 1.0. Ci < 1 was considered as drug synergism. (I) Flow cytometry analysis showing apoptosis of normal (N) HSPCs (n=3) treated with either DMSO, olaparib (400 nM), venetoclax (150 nM), or a combination of olaparib and venetoclax for 48 hr. Data represent average of two to three independent experiments. Statistics were calculated with Student’s t-test; error bars represent means ± SD. \**P* < 0.05 or \*\*\**P* < 0.001 were considered to be statistically significant.

Table S1. Details of AML patient sample characteristics.

Table S2. List of qRT-PCR primers.

Table S3. List of qChIP primers.

Dataset S1. List of differentially expressed genes in *Utx^+/+^* versus *Utx^-/-^* hematopoietic cells.

Dataset S2. List of Kdm6a ChIP-seq occupied genes in *Utx^+/+^*Lin^-^ bone marrow cells.

Dataset S3. List of genes with overlap for KDM6A and SMARCC1 occupancy.

Dataset S4. List of genes with overlap for KDM6A, SMARCC1 and H3K27ac occupancy while devoid of H3K27me3.

