## Supplementary Table S1 for "KDM6A loss sensitizes human acute myeloid leukemia to PARP and BCL2 inhibition"

**Table S1. Details of AML patient sample characteristics.**

| Samples | Disease type | KDM6A mutation type | Variant (cDNA) | Variant (AA) | Annotation |
|---|---|---|---|---|---|
| AML090147 | AML | Wild Type | -- | -- | -- |
| AML080527 | AML | Wild Type | -- | -- | -- |
| AML110002 | AML | Wild Type | -- | -- | -- |
| AML0646 | AML with intermediate cytogenetics, 48,XX,+8,+20[16], ITD negative | Nonsense | c.1347G>A | p.W449* | -- |
| AML208316 | AML with t(8;21)(q22;q22) | Frameshift | c.3336_3337insTCCTTTAACCC | p.Val1113Serfs*11 | Oncogenic |
| AML160326 | AML with MRC, complex karyotype including 17 p rearrangement, monosomy9, trisomy 1 | Frameshift | c.4188dupC | p.Thr1397Hisfs*2 | Oncogenic |
| AML188918 | AML with myelodysplasia-related changes | Missense | c.409G>T | p.Gly137Cys | TPR domain |
| AML161669 | AML with myelodysplasia-related changes/ 46, XX, del(5)(q13q33) | Missense | c.3293T>G | p.Leu1098Trp | Jmjc domain missense mutation |
| AML848978 | AML with mutated NPM1 | Frameshift | c.660dupA | p.Tyr221Ilefs*9 | Oncogenic |
