## Supplementary Table S2 for "KDM6A loss sensitizes human acute myeloid leukemia to PARP and BCL2 inhibition"

**Table S2. List of qRT-PCR primers.**

| Gene name | Primer sequence (5'-3') |
|---|---|
| *GAPDH_F* | GGAGTCCACTGGCGTCTTCAC |
| *GAPDH_R* | CTGATGATCTTGAGGCTGTTGTCA |
| *KDM6A_F* | CATGTTCCCTGTAGCACATCAAG |
| *KDM6A_R* | GGTCAGGTTTGTGCGGTTATG |
| *KDM6B_F* | AATCAGCGACCCCGACTTG |
| *KDM6B_R* | CATCGCACTCGTTGCAGTAGTAG |
| *BRCA1_F* | AGCATCTGGGTGTGAGAGTGAAAC |
| *BRCA1_R* | TCCTGCTGGAGCTTTATCAGGTTAT |
| *BRCA2_F* | TCTGCTGAACAAAAGGAACAAGGT |
| *BRCA2_R* | TCTGATGATGGACGCCAAATACTC |
| *RAD50_F* | ATACAAGCAACAAAATAGCACAGGATAAA |
| *RAD50_R* | TTGCTTCTTATAGTCGTCTTTCCCATCT |
| *RAD51_F* | CACCGCCCTTTACAGAACAGACTA |
| *RAD51_R* | TGATTACCACTGCTACACCAAACTCA |
| *RAD52_F* | CTAAATAAGCTTCCACGCCAGTTG |
| *RAD52_R* | GGTGTCCCAGGGCCATGTT |
| *XRCC2_F* | CAGAAAGCCTCGAGCTCATCAG |
| *XRCC2_R* | AAAAACATCCTGTGCTTCACCAGTT |
| *XRCC4_F* | GTACTGATGAGGAAAGTGAAAACCAAAC |
| *XRCC4_R* | TGTCTCCTTTTTCTACTTGGTGCAA |
| *XRCC5_F* | AGTAACCAGCTCATAAATCACATCGAA |
| *XRCC5_R* | CGCTGCTCTTCTGAAAACTTAATGG |
| *XRCC6_F* | GGAAGAAGAGTTGGATGACCAGAAAA |
| *XRCC6_R* | CTGCTCTGGAGTTGCCATGATT |
| *LIG4_F* | ATGATTCATATGTGCCCATCAACCA |
| *LIG4_R* | CATTTCTTCAGGAGTCTGCTCGTTAG |
| *BCL2_F* | CATGTGTGTGGAGAGCGTCAAC |
| *BCL2_R* | GGGCCGTACAGTTCCACAAAG |
| *BCL-xL_F* | CAGAAGGGACTGAATCGGAGATG |
| *BCL-xL_R* | CTCCCTCAGCGCTTGCTTTAC |
| *MCL1_F* | CACCAGTACGGACGGGTCAC |
| *MCL1_R* | GCCCCAGACCTGCCCATT |
| *BCL2A1_F* | AGAACACTATTCAACCAAGTGATGGA |
| *BCL2A1_R* | TTGCTGTCGTAGAAGTTTCTTGATG |
| *HOXA9_F* | GGCGGTGCCCCTATACAAAAC |
| *HOXA9_R* | CTTCATTTTCATCCTGCGGTTCTG |
| *MRE11A_F* | TCAAGATGGCAACCTCAACATTTC |
| *MRE11A_R* | TTGAAGCAAAACCGGACTAATGTC |
| *ATM_F* | CTCTCAGCGAAGTGGTGTTCTTG |
| *ATM_R* | TTGCACCTCCATCATTTTCTTTTG |
