## Supplementary Table S3 for "KDM6A loss sensitizes human acute myeloid leukemia to PARP and BCL2 inhibition"

**Table S3. List of qChIP primers.**

| Gene name | Primer sequence (5'-3') |
|---|---|
| *KDM6A_TSS_F* | TGACGTGAGTCAACAAAGGTCAC |
| *KDM6A_TSS_R* | GCACCAGCACAAAGTTGTTGTAA |
| *KDM6B_TSS_F* | ACGGGCAACCCCCGAAAT |
| *KDM6B_TSS_R* | CCGACCCCCAAAATCAGACC |
| *BRCA1_TSS_F* | ACGGAAAAGCGCGGGAATTA |
| *BRCA1_TSS_R* | TACCCAGAGCAGAGGGTGAA |
| *BRCA1_U1_F* | TACGAAAACATAACACTCCAGTCCA |
| *BRCA1_U1_R* | TCTTCGCCACAGTGTTCCTTA |
| *BRCA2_TSS_F* | CTGAAACTAGGCGGCAGAGG |
| *BRCA2_TSS_R* | GGCCGGAGTAAGCTGACAAA |
| *BRCA2_U1_F* | GCGAGTTAAGATGGGTTTCACAA |
| *BRCA2_U1_R* | AGTGGGCGGGGCTGTTATT |
| *RAD50_TSS_F* | GGACTCTGATTTCCCGGCGT |
| *RAD50_TSS_R* | AAGCCGTAGCCACAATGCG |
| *RAD50_U1_F* | CCCTCCATTAAGTTTGTTTCCAAGA |
| *RAD50_U1_R* | GGTAATCCATTCCGATCAGTGC |
| *RAD51_TSS_F* | TCTTGGGTTAGCGCGCAG |
| *RAD51_TSS_R* | ACTCTCCTTAGGGCTCGGT |
| *RAD51_U1_F* | CCCGTACGCTAGCTCCATT |
| *RAD51_U1_R* | TTTGGCACTTCTGGTCGCT |
| *RAD52_TSS_F* | CCCCTCCGACTTGATTTACGG |
| *RAD52_TSS_R* | AGGGAGCTCGATCTAGGCTAT |
| *RAD52_U1_F* | GGGGACATTCTAGAGGGGACT |
| *RAD52_U1_R* | GTTCTCGCGGCCTCTCAC |
| *BCL2_ TSS_ F* | AACCGGTCGGGCTGTGC |
| *BCL2 _TSS_ R* | ACCTTCGCTGGCAGCG |
| *BCL2_U1_F* | GTTCCCGGCGCTTTTCCAG |
| *BCL2_U1_R* | GCGGACTTGGTGGTCGCT |
| *BCL2A1_TSS_F* | ACACAGCCTACGCACGAAA |
| *BCL2A1_TSS_R* | GAGCAAAGTCTTGAGCTGGC |
| *BCL2A1_U1_F* | GCGGCCGGCCTTGTAAA |
| *BCL2A1_U1_R* | AAATCCATAGGATAGAAAGCAAATTGG |
